# Evidence for a role of phenotypic mutations in virus adaptation

**DOI:** 10.1101/2020.08.21.261495

**Authors:** Raquel Luzon-Hidalgo, Valeria A. Risso, Asuncion Delgado, Eduardo Andrés-Leon, Beatriz Ibarra-Molero, Jose M. Sanchez-Ruiz

## Abstract

Viruses interact extensively with the host molecular machinery, but the underlying mechanisms are poorly understood. Bacteriophage T7 recruits the small protein thioredoxin of the *E. coli* host as an essential processivity factor for the viral DNA polymerase. We challenged the phage to propagate in a host in which thioredoxin had been extensively modified to hamper its recruitment. The virus adapted to the engineered host without losing the capability to propagate in the original host, but no genetic mutations were fixed in the thioredoxin binding domain of the viral DNA polymerase. Virus adaptation correlated with mutations in the viral RNA polymerase, supporting that promiscuous thioredoxin recruitment was enabled by phenotypic mutations caused by transcription errors. These results point to a hitherto unknown mechanism of virus adaptation that may play a role in crossspecies transmission. We propose that phenotypic mutations may generally contribute to the capability of viruses to evade antiviral strategies.

## INTRODUCTION

Viruses repurpose the host molecular machinery for their own proliferation, block or evade antiviral factors and recruit host proteins (proviral factors) for processes critical for virus propagation. Viruses occasionally jump between species, sometimes with fatal consequences for the new host. Homolog proteins that carry out the same tasks in two different organisms likely share functionality and 3D-structure, but differ in amino acid sequence and, consequently, they also differ in the chemical composition of their exposed surfaces available for intermolecular interaction. In view of this, crossspecies transmission is quite remarkable, as it implies that the virus can simultaneously establish extensive and effective interactions in the two quite different molecular environments of the new and the old hosts. To explore fundamental mechanisms behind this molecular-level promiscuity, we have carried out laboratory evolution experiments in which a lytic phage is challenged to propagate in an engineered host that had been modified to hinder the recruitment of a known proviral factor. Our experiments target a specific virus-host interaction for which extensive functional and structural information is available, thus considerably facilitating the interpretation of the experimental results. Indeed, while we observe quick and efficient evolutionary adaptation of the virus to the engineered host, DNA sequencing reveals that no inheritable, genetic mutations occur at the targeted virus-host interaction. This essentially leaves phenotypic mutations caused by transcription errors as the only possible mechanism of adaptation in this case, a scenario which is actually consistent with the mutational pattern we find in the viral RNA polymerase.

Mistakes in protein synthesis, due to translation and transcription errors, are common and lead to the so-called phenotypic mutations (Drummond and Wilke, 2009; Goldsmith and Tawfik, 2009). Translation error rates are generally higher than transcription error rates. Yet, transcription errors may have a stronger impact at the protein level, since an mRNA molecule is typically translated many times (Traverse and Ochman, 2016). Regardless of their origin, phenotypic mutations are not inherited and are often regarded as a burden to the organism because they may result in a substantial amount of misfolded protein molecules that are non-functional and that may actually be harmful (Drummond and Wilke, 2009; Goldsmith and Tawfik, 2009). On the other hand, some phenotypic mutations could provide crucial functional advantages under certain conditions. It has been theorized that phenotypic mutations may play an evolutionary role by allowing organism survival until functionally useful mutations appear at the genetic level (Whitehead et al., 2008; Petrovic et al., 2018). The laboratory evolution experiments reported here support this hypothesis while providing evidence for a hitherto unknown mechanism of virus adaptation.

More generally, we argue here that an adaptation mechanism based on phenotypic mutations has important implications beyond the specific model system used here to demonstrate it. Many key intermolecular interactions may need to occur only a few times per host cell to allow virus propagation and can, therefore, be realized by mutant proteins present at very low levels. It follows that diversity at the phenotypic level caused by transcription errors may generate an extensive capability to establish/avoid interactions in different molecular environments, thus facilitating crossspecies transmission and potentially contributing to the capability of viruses to evade antiviral strategies.

## RESULTS

### Design of the engineered host

Bacteriophage T7 recruits the small (~110 residues) protein thioredoxin from the *E. coli* host to be a part of its four-protein replisome (Hamdan and Richardson, 2009). Upon recruitment, thioredoxin becomes an essential processivity factor for the viral DNA polymerase (Etson et al., 2010) (Figure 1A). In the absence of thioredoxin, the processivity of the DNA polymerase is only 1-15 nucleotides per binding event (versus several hundred upon thioredoxin binding), a very low level which does not lead to virus replication and propagation in the knockout strains used in this work (see below). Binding of *E. coli* thioredoxin to the viral DNA polymerase is mediated by the interaction with a unique 76-residue fragment often referred to as the thioredoxin binding domain or TBD (Hamdan and Richardson, 2009; Akabayov et al., 2010; Lee and Richardson, 2011). This interaction is extremely tight (dissociation constant ~5 nanomolar) showing that the TBD has evolved to specifically bind the host thioredoxin (Hamdan and Richardson, 2009). It follows that replacing *E. coli* thioredoxin with an alternative thioredoxin could hinder recruitment and potentially prevent phage propagation in *E. coli* (Delgado et al., 2017). A simple way to perform such replacement involves complementing a knockout *E. coli* Trx^−^ strain with a plasmid bearing the alternative thioredoxin gene (Delgado et al., 2017). The engineered host used in the experiments reported here is further modified to allow phenotypic mutations linked to viral transcription errors to occur in both partners of the targeted DNA polymerase-thioredoxin interaction. That is, the DNA polymerase is transcribed by the viral T7 RNA polymerase and, in addition, we placed the alternative thioredoxin gene in the plasmid under a promoter of the viral RNA polymerase. The gene for the wild type T7 RNA polymerase was inserted in the host chromosome, which ensures the presence of a significant amount of thioredoxin even before phage infection. In order to have available a suitable control host, the same procedure was performed with an *E. coli* Trx^−^ complemented with a plasmid bearing the gene of *E. coli* thioredoxin. The resulting strain is similar to *E. coli* in terms of growth and susceptibility to bacteriophage T7 (Delgado et al., 2017) and has been used as a representation of the original host throughout this work.

**Figure 1.**
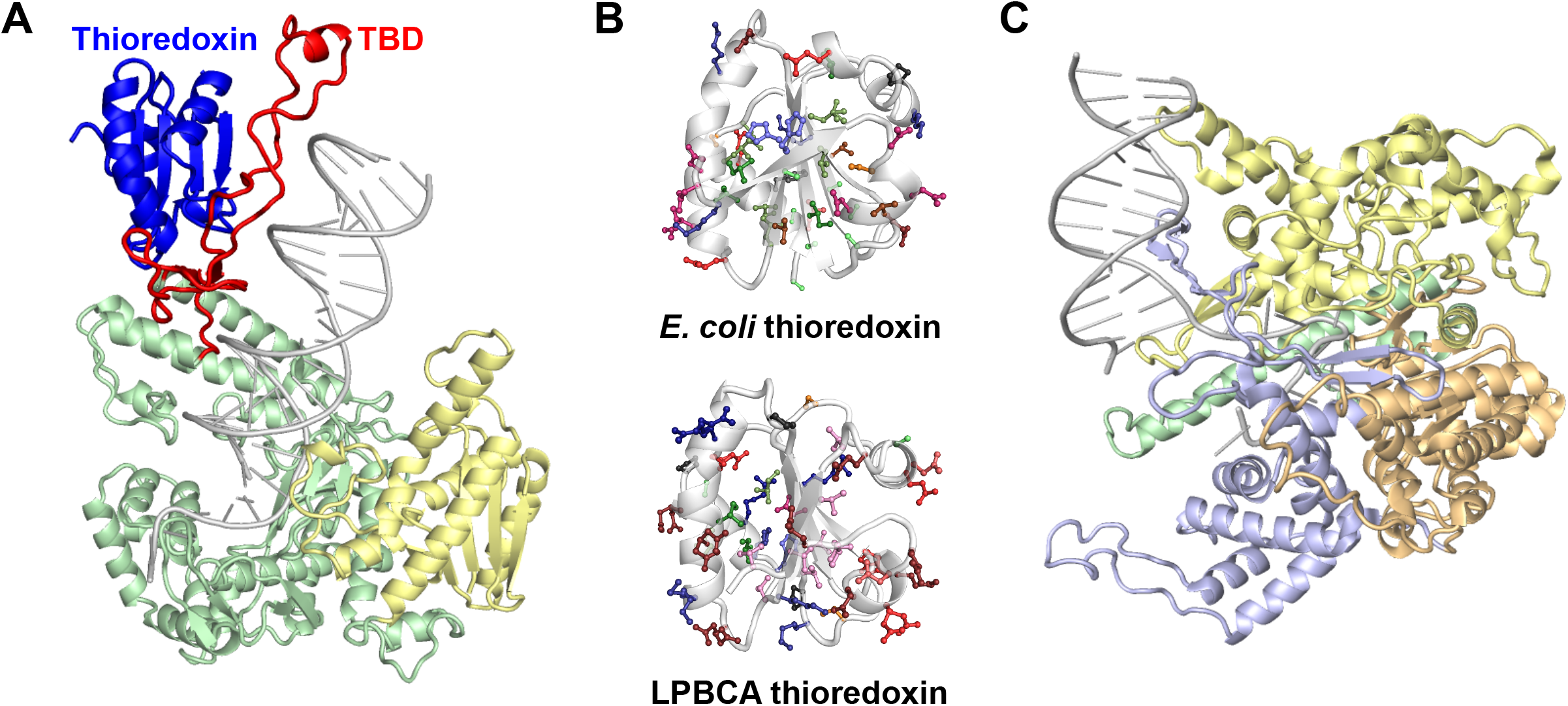
3D structures of the main molecular players in this work. **A,** Structure of the viral DNA polymerase interacting with its processivity factor, *E. coli* thioredoxin (PDB ID 1T8E). The thioredoxin binding domain of the viral polymerase is labelled with TBD. **B,** Structures of *E. coli* thioredoxin (PDB ID 2TRX) and the alternative LPBCA thioredoxin (PDB ID 2YJ7) used in this work. The amino acid residues that differ between the two structures are shown in stick representation. **C,** Structure of the viral RNA polymerase (PDB ID 1QLN).

As alternative thioredoxin we used a protein, LPBCA thioredoxin, previously obtained and characterized in detail as part of an ancestral reconstruction study (Perez-Jimenez et al., 2011; Ingles-Prieto et al., 2013). Actually, LPBCA stands for “last common ancestor of cyanobacterial, deinococcus and thermos groups”. LPBCA thioredoxin shares function and 3D-structure with *E. coli* thioredoxin, but displays only 57% sequence identity with its modern counterpart. As a result, the amino acid composition of its exposed protein surfaces is substantially altered with respect to *E. coli* thioredoxin (Figure 1B) and efficient interaction with the thioredoxin binding domain of the viral DNA polymerase is not likely to occur (Delgado et al., 2017). The putative ancestral nature of LPBCA thioredoxin is, of course, immaterial for this work. The key features of LPBCA thioredoxin in the context of this work are that 1) it poses an *a priori* tough challenge to the virus and 2) we know beforehand that mutations enabling recruitment of the alternative thioredoxin must be at the TBD-thioredoxin interaction surface. This fact facilitates considerably the interpretation of the experimental data. Figure 2 shows schematically the *E. coli* variants used in this work as well as their main features as hosts for phage replication.

**Figure 2.**
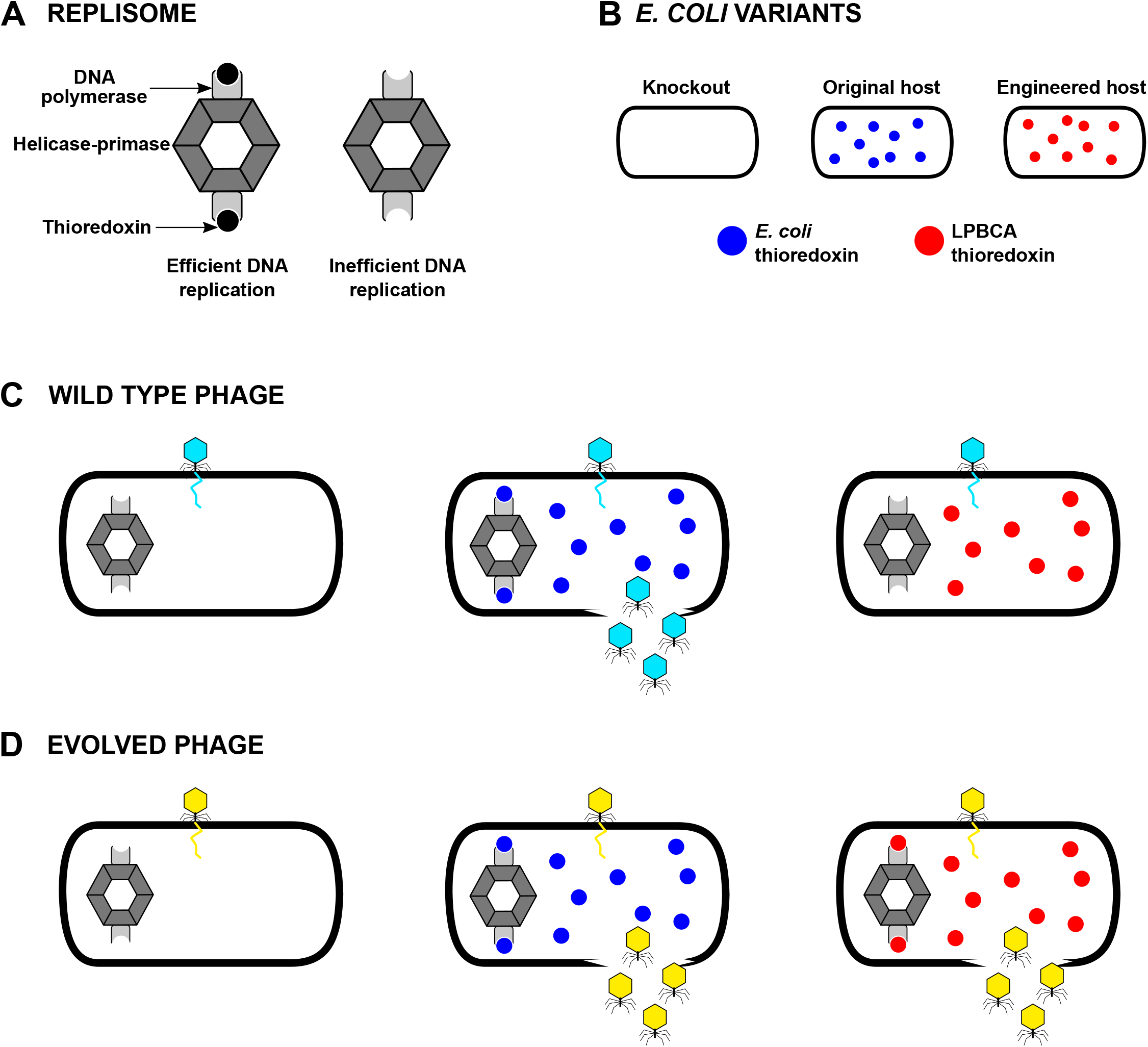
*E. coli* variants used in this work and their behaviour as hosts for phage replication. **A,** *E. coli* thioredoxin is recruited for the T7 bacteriophage replisome as an essential processivity factor for the viral DNA polymerase. A cartoon representation of the replisome is shown here. See figure 1 in Hamdan and Richardson (2009) for a more realistic model. **B,** Differences between the three *E. coli* variants used in this work are related to thioredoxin. An *E. coli* trx-knockout strain lacks thioredoxin. A control strain expresses the “normal” *E. coli* thioredoxin and it is used as our representation of the original host. The engineered host expresses the alternative LPBCA thioredoxin, which displays only 57% sequence identity with *E. coli* thioredoxin (see Figure 1B). **C,** DNA replication for wild-type T7 bacteriophage requires recruitment of *E. coli* thioredoxin and efficient virus propagation is only observed with the original host. **D,** Results reported in this work show, however, that the virus can evolve to propagate in the engineered host. The evolved virus does not propagate in the knockout host, but still propagates efficiently in the original host, pointing to a promiscuous capability to recruit both *E. coli* thioredoxin and LPBCA thioredoxin, as symbolically depicted by the replisomes shown.

### General design of the laboratory evolution experiments

In plaque assays with the *E. coli* Trx^−^ strain complemented with LPBCA thioredoxin (*i.e*., our engineered host), plaques are only observed at the highest virus concentrations used and, even under those conditions, they are observed only occasionally (Delgado et al., 2017). It appears then that only a tiny fraction of the virions in the phage sample (~10^−7^ or less) can actually propagate in the engineered host. Furthermore, the propagation is initially inefficient, as judged by the very small size of the plaques they generate (Figure S1). Here, we have evolved such “anomalous” virions for efficient propagation in the engineered host. Schematically, our evolution experiments can be represented as:

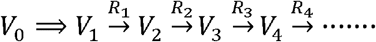

where V_0_ is the original virus sample, V_1_ is a virus sample obtained from the initial propagation in the engineered host (double-line arrow) and the V_2_, V_3_, etc. samples result from the R_1_, R_2_, etc. evolutionary rounds, each involving plaque assays for propagation in the engineered host with ten-fold dilutions and plaque selection (see Methods for details and Figure 3 for a pictorial representation of the evolution experiments). These assays immediately lead to numbers of virus particles (plaque forming units or pfu) that could infect the engineered host. For each plaque selected from the rounds of adaptation to the engineered host, we also performed plaque assays to determine the number of particles that could infect the original host.

**Figure 3.**
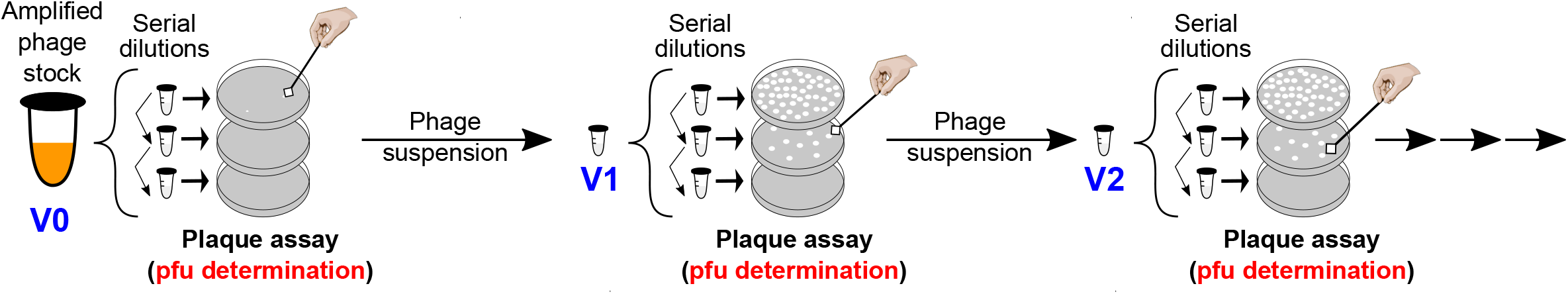
Initial phage evolution experiments performed in this work. In each evolutionary round, plaque assays with serial dilutions of a phage suspension are performed and, therefore, number of plaque forming units in the suspension can be calculated. At each round, a well-defined plaque from the highest dilution at which plaques are observed is picked and suspended in buffer to yield the phage suspension used to start the next round (see main text and Methods for further details). This protocol was used in the two long evolution experiments that led to pfu profiles of Figure 4 below.

### Evolution experiments reveal promiscuous viral adaptation

We first performed two long evolution experiments consisting of an evolutionary round per day during several weeks. In these experiments, samples corresponding to a plaque surface of 3×3 mm^2^ were used to start each next round (except for the V_1_ sample for which the whole plaque was used, since it was smaller than 3×3 mm^2^). Both experiments (Figure 4) revealed a fast adaptation of the virus to engineered host, as shown by increases in plaque size and in the numbers of plaque forming units (see Figure S2 for a representative example of the plaque size increases observed in this work). In fact, the numbers of plaque forming units determined using the engineered and the original host became similar to each other within the first 1-2 rounds and, surprisingly, the two numbers remained similar over many rounds of evolution. The two experiments were stopped at some point and restarted after about one month from plates stored at 4 °C. In one case (experiment in lower panel of Fig. 4), the protocol was changed after re-starting by eliminating an amplification step (see legend of Figure 4 for details). As expected, these one-month breaks are reflected in sudden drops in pfu numbers. Nevertheless, the pfu numbers determined using the original and the engineered hosts still remained very similar to each other. The congruence between the two pfu numbers is quite remarkable because we challenged the virus to propagate in the engineered host and we did not impose any selective pressure for propagation in the original host. Therefore, the capability of the evolved virus to propagate in the two hosts cannot be explained by assuming two genetically differentiated virus sub-species in our samples, because the subspecies competent to propagate in the engineered host would have quickly dominated the virus population in the evolution experiments. Obviously, whatever the virus adaptation mechanism is, it is intrinsically promiscuous, meaning that each infective virion can propagate in both hosts.

**Figure 4.**
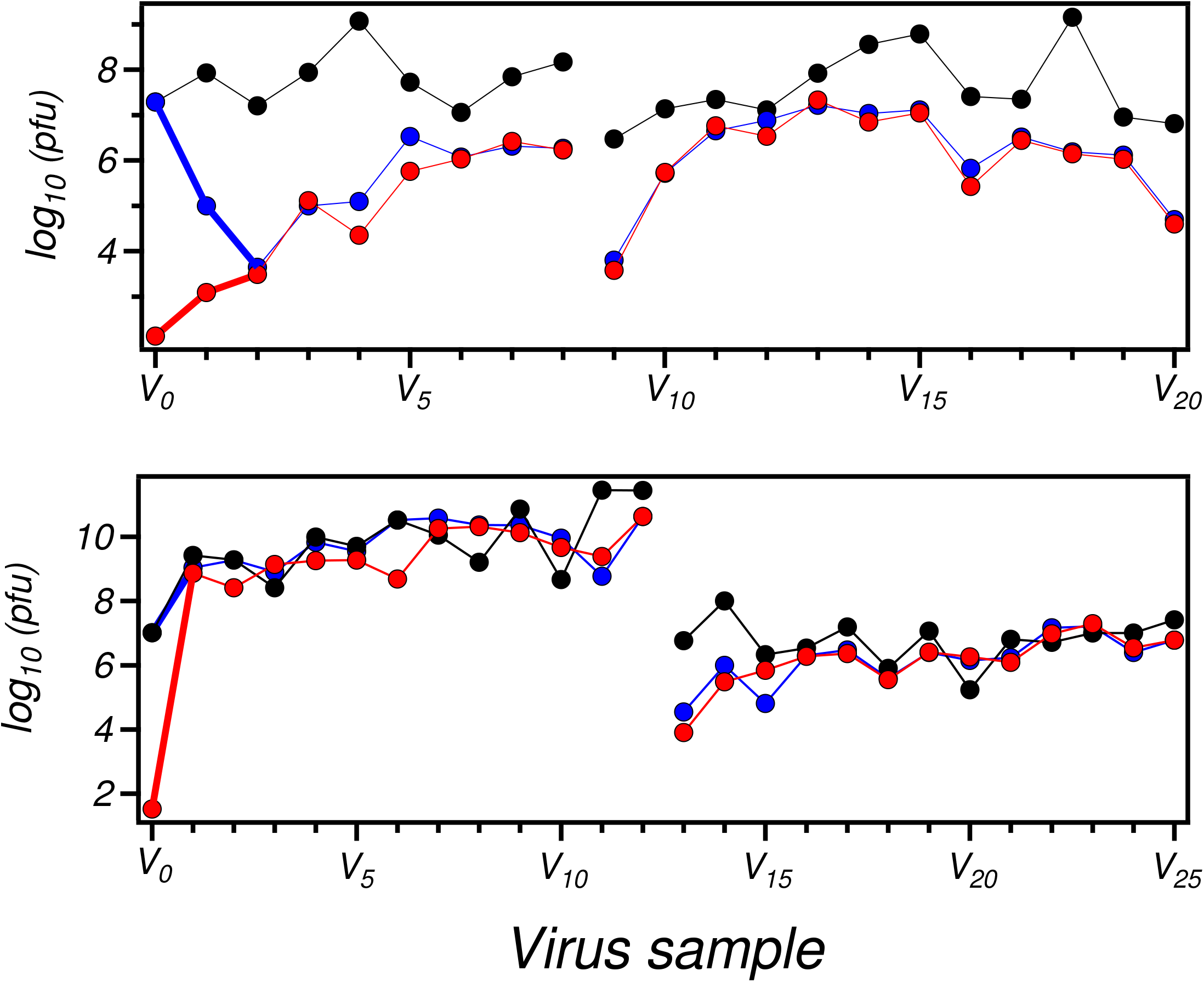
Propagation of bacteriophage T7 in engineered *E. coli* cells with a modified thioredoxin gene. Number of plaque forming units (pfu) for virus samples from two laboratory evolution experiments performed as described in the text. Pfu values determined using the engineered host (red) and the original host (blue) are shown. It is important to note, however, that only selection for propagation in the engineered host has been applied in these experiments. Data from control experiments are also shown (black). These control experiments used the original (non-evolved) phage and the original *E. coli* host and were carried out concurrently with the evolution experiments. Experiments were interrupted after a substantial number of rounds and restarted from stored plaques after about a month and continued for an additional number of rounds. The interruption is apparent from the break in the lines that connect the experimental data points.

### Virus adaptation and host promiscuity are not explained by genetic mutations

It is important to note that none of the many evolved virus samples studied in this work was found to propagate in the knock-out *E. coli* Trx^−^ strain that lacks thioredoxins. Therefore, the observed host promiscuity is not due to the virus evolving a capability to assemble a functional replisome without the assistance of thioredoxin. Rather, it must be linked to a capability of the viral DNA polymerase to recruit the thioredoxins in both the original and the engineered hosts. In order to determine the mutations in the viral DNA polymerase gene that could potentially be responsible for the promiscuous recruitment, we carried out 14 “short” evolution experiments and we performed PCR followed by Sanger sequencing on samples from the early rounds (see Figure 5 for a pictorial representation of these evolution experiments). Between 1 and 3 genetic mutations appeared in the DNA polymerase in most experiments (see Figure 6 and also Table S1 for details), but none of them occurred in the thioredoxin binding domain. Interestingly, however, the 21 positions at which mutations are found (including data from the {A,B,C} experiments discussed below) define a clear structural pattern (Figure 6), clustering in the region of the polymerase domain close to the bound DNA and in the exonuclease domain involved in proofreading. Therefore, none of the mutations found can reasonably explain thioredoxin recruitment in the engineered host and, certainly, they do not allow efficient replication in the absence of thioredoxin, since the evolved virus does not propagate in the knock-out *E. coli* Trx^−^ strain. However, it is plausible that these mutations increase replication errors, thus promoting mutations in other viral proteins that trigger processes that eventually lead to recruitment. This interpretation is supported by experiments and analyses given further below.

**Figure 5.**
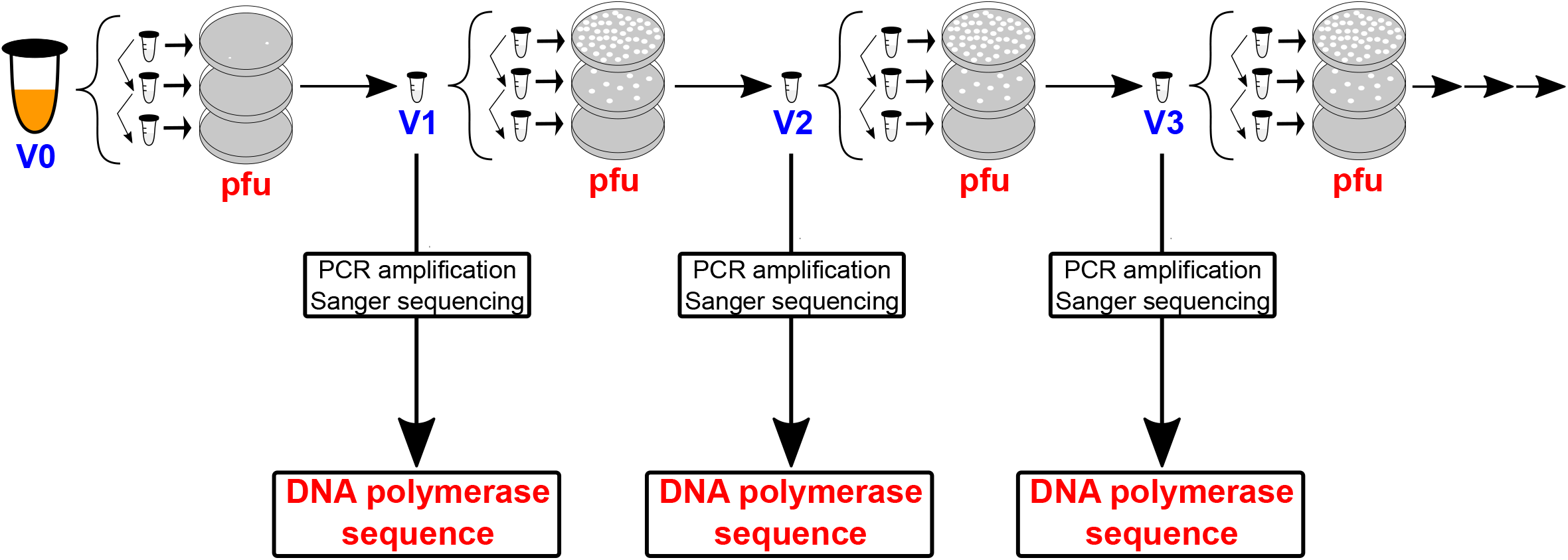
Evolution experiments modified to include DNA polymerase sequencing. The protocol described in Figure 3 can be easily modified to include PCR amplification and Sanger sequencing of the DNA polymerase for each phage suspension. This modified protocol was used in the 14 short evolution experiments that led to the sequence information on the viral DNA polymerase shown in Figure 6 and Table S1.

**Figure 6.**
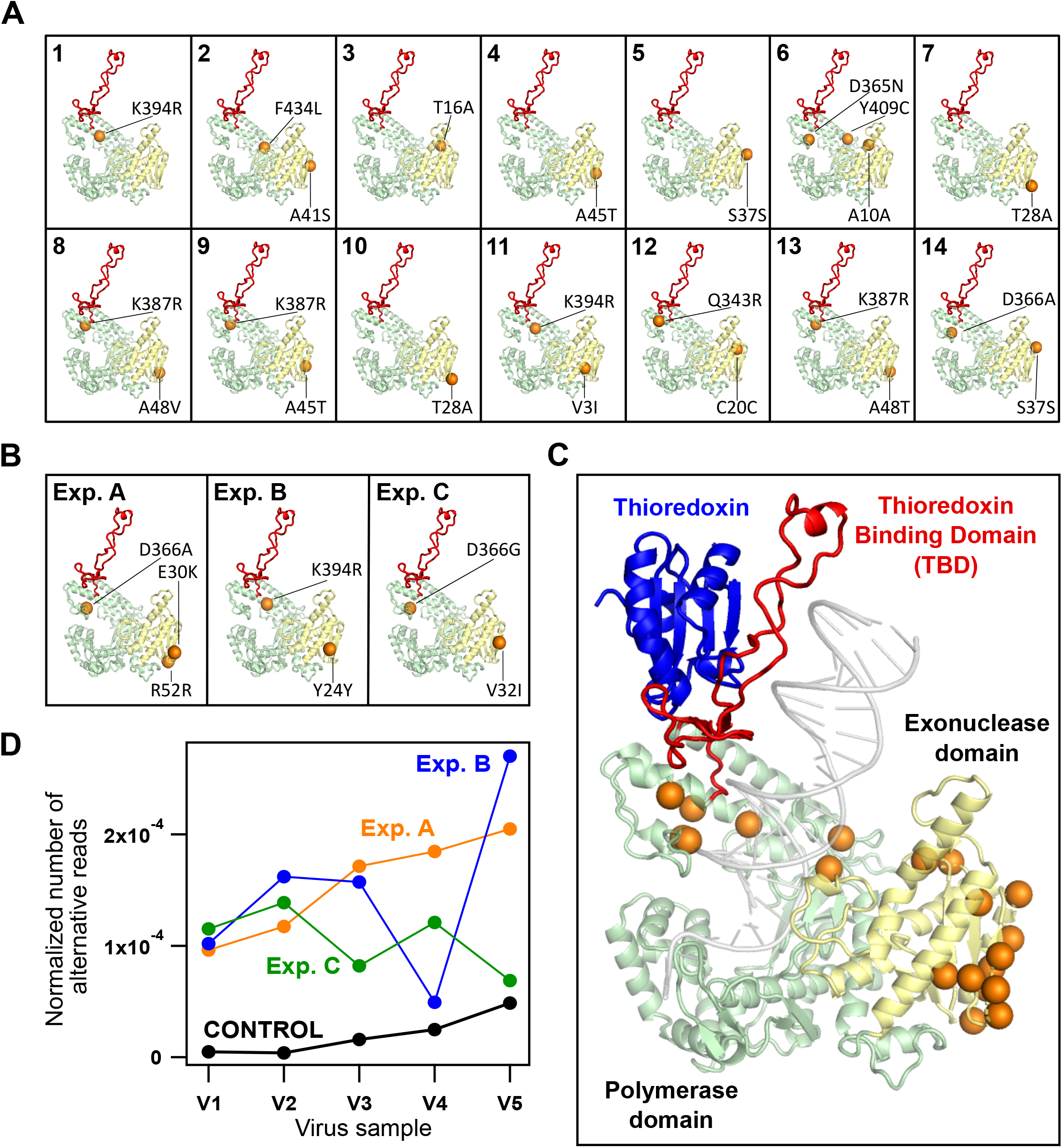
Structure of the viral DNA polymerase showing mutations appearing during phage adaptation to the engineered host. **A,** Mutations found in 14 “short” evolution experiments consisting of 2-3 rounds after PCR and Sanger sequencing of the DNA polymerase (see main text, Figure 5 and Table S1 for details). **B,** Mutations observed at high frequency (fraction > 0.95) in the {A,B,C} set of evolution experiments, as determined from next generation sequencing of the viral genome (see Figure 7 and main text for details). The mutations shown are those determined for the virus samples obtained after the fourth round (V5 samples). See Table S2, Figure 9 below and Figure S3 for the mutations in all viral samples during the evolution experiment. **C,** All sites at which mutations were found are highlighted with spheres in a single DNA polymerase structure. It is visually apparent that no mutations were found in the thioredoxin binding domain (TBD). **D,** Plot of normalized number of alternative reads in the genome of virus samples from the several rounds of the {A,B,C} experiments compared with a control experiment in which the non-evolved phage was allowed to propagate in the original host. These values support that the mutations fixed in the DNA polymerase during the {A,B,C} experiments (panel **B**) increase the replication errors (see text for details).

### Virus adaptation probed by lysis profiles

Host promiscuity, as described in the preceding sections, was inferred from the congruence between the numbers of plaque forming units determined using the engineered host and the control, “original” host. The implication is, therefore, that each infective virion particle from the evolved virus sample is capable of replicating in both hosts. This, however, does not necessarily mean that the efficiency of virus replication is the same in the two hosts. In order to investigate replication efficiency, we performed three additional evolution experiments using a modified protocol that allows lysis profiles to be determined (see Figure 7 for a pictorial representation of these experiments). We label these experiments as A, B and C and collectively refer to them as to the {A,B,C} set. Briefly, experiments were performed as described above, but, in addition, aliquots of the virus suspension at each round were used to assess lysis in solution with the original host, the engineered host and, as a control, with the knockout *E. coli* Trx^−^ strain.

**Figure 7.**
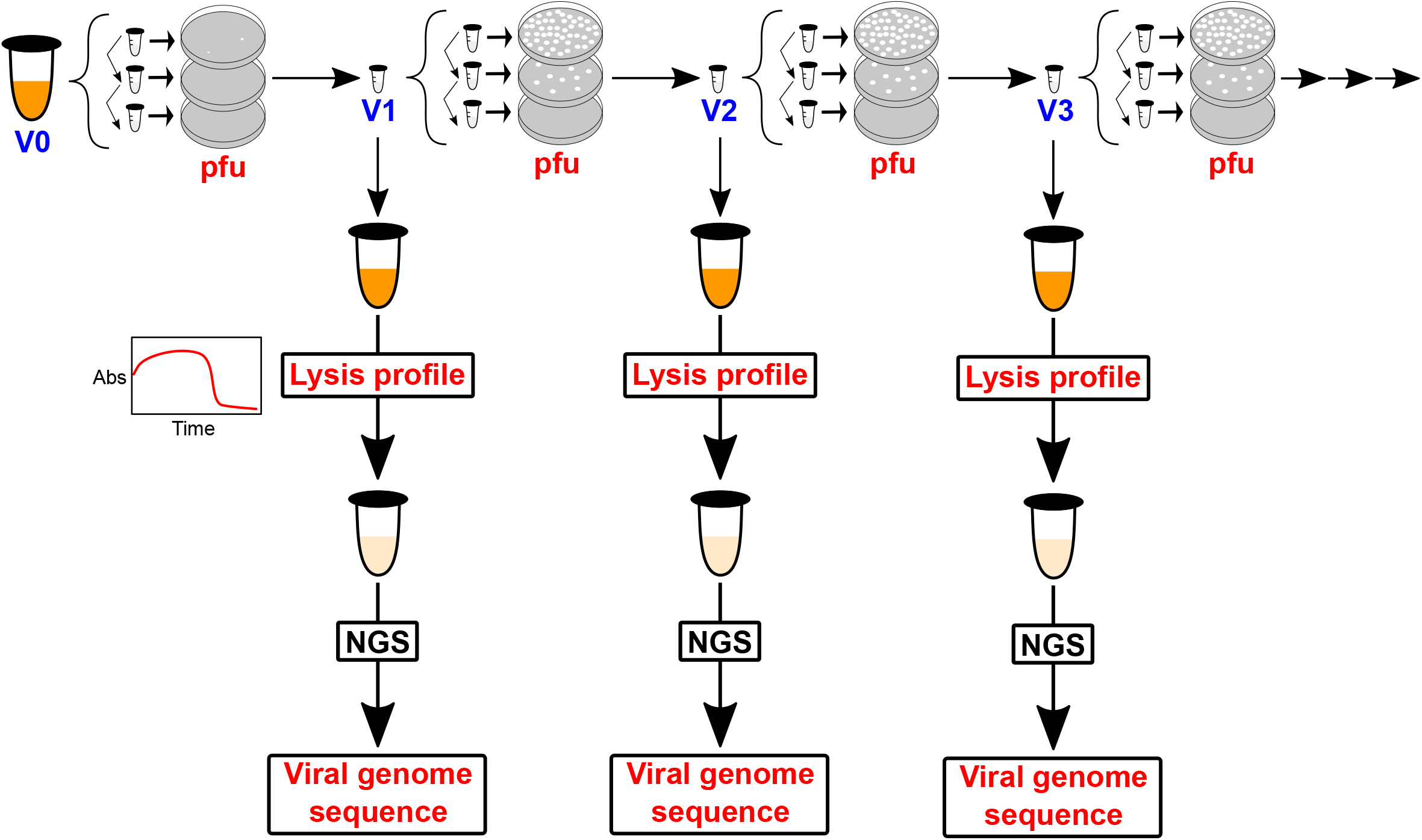
Evolution experiments modified to include the determination of lysis curves and NGS of the viral genome. The evolution experiments described in Figure 3 can be easily modified to determine *E. coli* lysis profiles (turbidity *versus* time) for the phage from the suspensions at each evolutionary round (see Methods for details). Furthermore, DNA extraction after lysis allows next generation sequencing of the viral genome at each evolutionary round (see Methods for details). This modified protocol was used in the {A,B,C} set of evolution experiments that led to the lysis profiles of figure 8 and the sequence information provided in Figures 9, 10, S3-S7 and Table S2.

In the three experiments of the {A,B,C} set, the phage adapted to the engineered host similarly to what was observed in the previous evolution experiments discussed above. Adaptation was thus immediately clear from the general trend towards increased plaque sizes in the first rounds. It is to be noted, however, that, in order to maximize the amount of sample for DNA extraction, the largest plaque was *fully* removed from the agar plate with the smaller number of plaques to start each next round in the {A,B,C} experiments. This leads to large increases in the determined pfu numbers over the evolution experiment. Actually, the combined pfu data for the three experiments span 5 orders of magnitude (Figure 8A). Still, there is clear congruence between the pfu values determined using the engineered and the original hosts over this very wide range, showing again that the mechanism of virus adaptation is intrinsically promiscuous.

**Figure 8.**
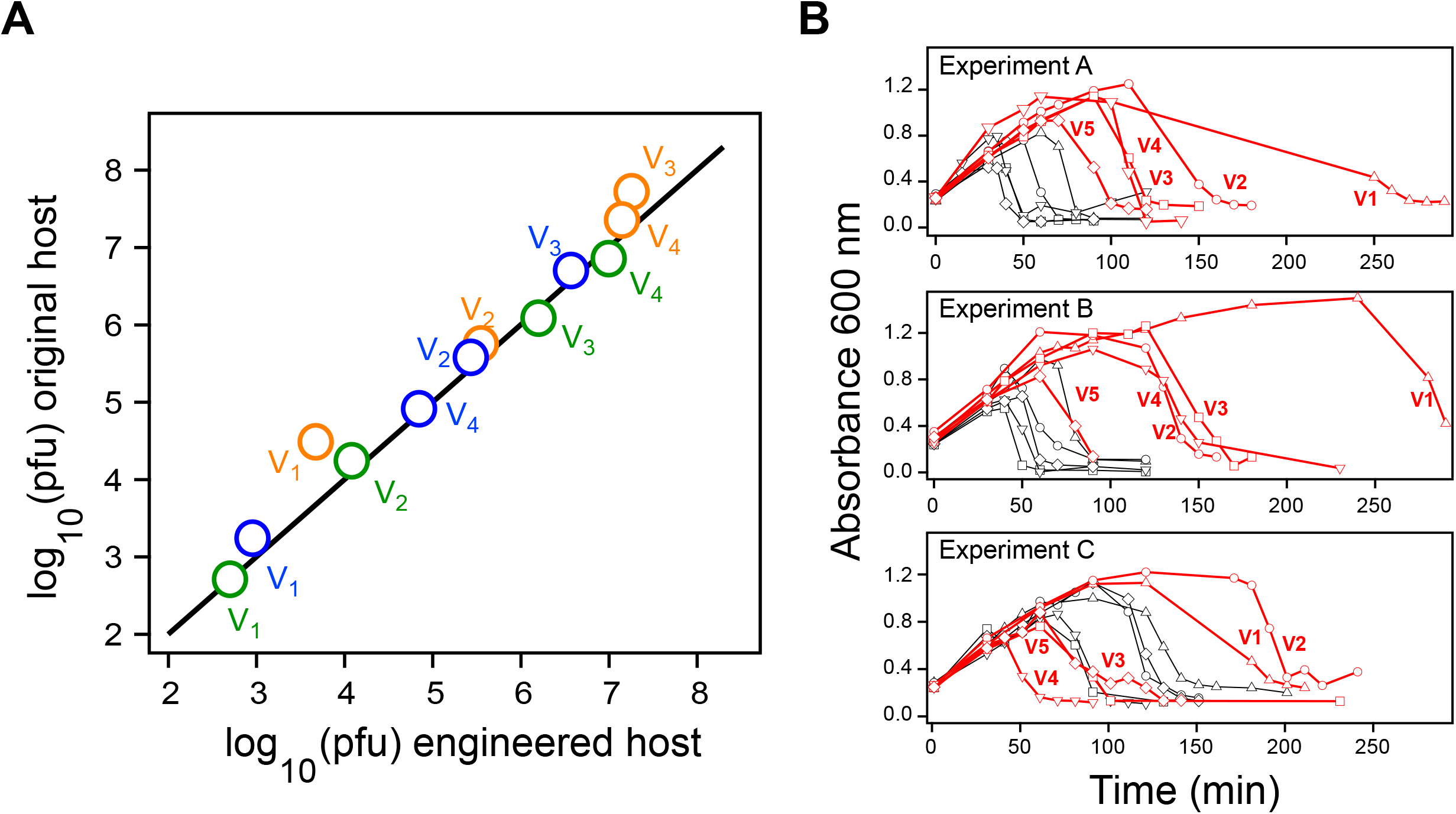
Engineered host versus original host adaptation during laboratory virus evolution. Data from the {A,B,C} set of laboratory evolution experiments (see text for details). The three experiments involve selection for propagation in the engineered host. **A,** Pfu values for virus samples from the evolution experiments determined with the original host versus the values determined with the engineered host. Note that the values span about 5 orders of magnitude. The line is not a fit but represents the equality of the two pfu values. Color code identifies the experiment: A(orange), B (blue), C (green). **B,** Lysis plots of absorbance at 600 nm versus time for the virus samples from the three evolution experiments. Absorbance at 600 nm reflects turbidity and lysis is revealed by an absorbance drop. Profiles determined using the engineered host (red) and the original host (black) are shown. For both the original host and the engineered host, symbols identify the virus sample: triangles (V_1_), circles (V_2_), squares (V_3_), down pointing triangles (V_4_), diamonds (V_5_). Lysis was not observed in control experiments with the knockout *E. coli* Trx^−^ strain that lacks thioredoxin.

Lysis profiles for experiments of the {A,B,C} set reveal that, although infective virions can replicate in the original and the engineered hosts, they certainly do not do so with the same efficiency, at least in the first evolutionary rounds. As shown in Figure 8B, lysis times decrease over the evolution rounds reflecting, at least in part, the increases in viral load associated with the use of full plaques to start rounds. More relevant is the fact that, initially, lysis times are clearly larger for the engineered host as compared with the original host (Figure 8B). Upon virus adaptation, however, lysis times for the engineered host approach those for the original host, although to an extent that it is variable. Lysis times for the engineered host always remain larger in one experiment (labelled A), while they eventually become somewhat smaller than the lysis times for the original host in another experiment (labelled C). As expected, no lysis was observed with the knockout *E. coli* Trx^−^ strain that lacks thioredoxin, not even after overnight incubation. Overall, from the point of view of engineered versus original host replication efficiency, the experiments in Figure 8B can be ranked C>B>A, with the later virus samples from experiment C being the most efficient at replicating in the engineered host.

### Next generation sequencing studies on virus adaptation to the engineered host

The protocol used for the {A,B,C] experiments allows sequencing of the viral genome at each evolutionary round (see Figure 7). Thus, DNA was extracted from the lysed engineered host samples and used for Illumina next generation sequencing. Certainly, lysis experiments involve an additional evolution step in solution that can potentially result in additional mutations. Still, as it will be apparent from the discussions further below, the DNA sequencing information from the lysed samples provides a clear picture of the molecular mechanism behind viral adaptation.

The Illumina sequencing data for the {A,B,C} set were processed in two ways. First, single nucleotide variants (SNVs) that appeared at high frequency (fraction over the total of readings at the position >0.95) were determined for all viral genes. These results are shown in Figures 9 and S7 and highlighted in yellow in Table S2. Secondly, for genes of particular interest, we considered all SNVs fulfilling these two criteria: 1) the fraction of the nucleotide variant at a given position over the total number of readings was 0.01 or higher; 2) the SNV is observed at least two times. Overall, for both, the DNA polymerase and the RNA polymerase, the selected SNVs define clearly bimodal distributions, including low frequency SNVs (fraction over total number of readings between 0.01 and 0.05) and high frequency SNVs (fraction between 0.95 and 1) with no SNVs at intermediate fractions in most cases (Figures S3 and S4). The numbers of low frequency SNVs are shown in Figure 10, while the mutations themselves are given in Table S2. Note that most of these low-frequency SNVs appear only transiently, are not enriched and do not become fixed. On the other hand, for most of the mutations that do become fixed (see files highlighted in yellow in Table S2), the mutation is often detected at a lower level in an early round.

**Figure 9.**
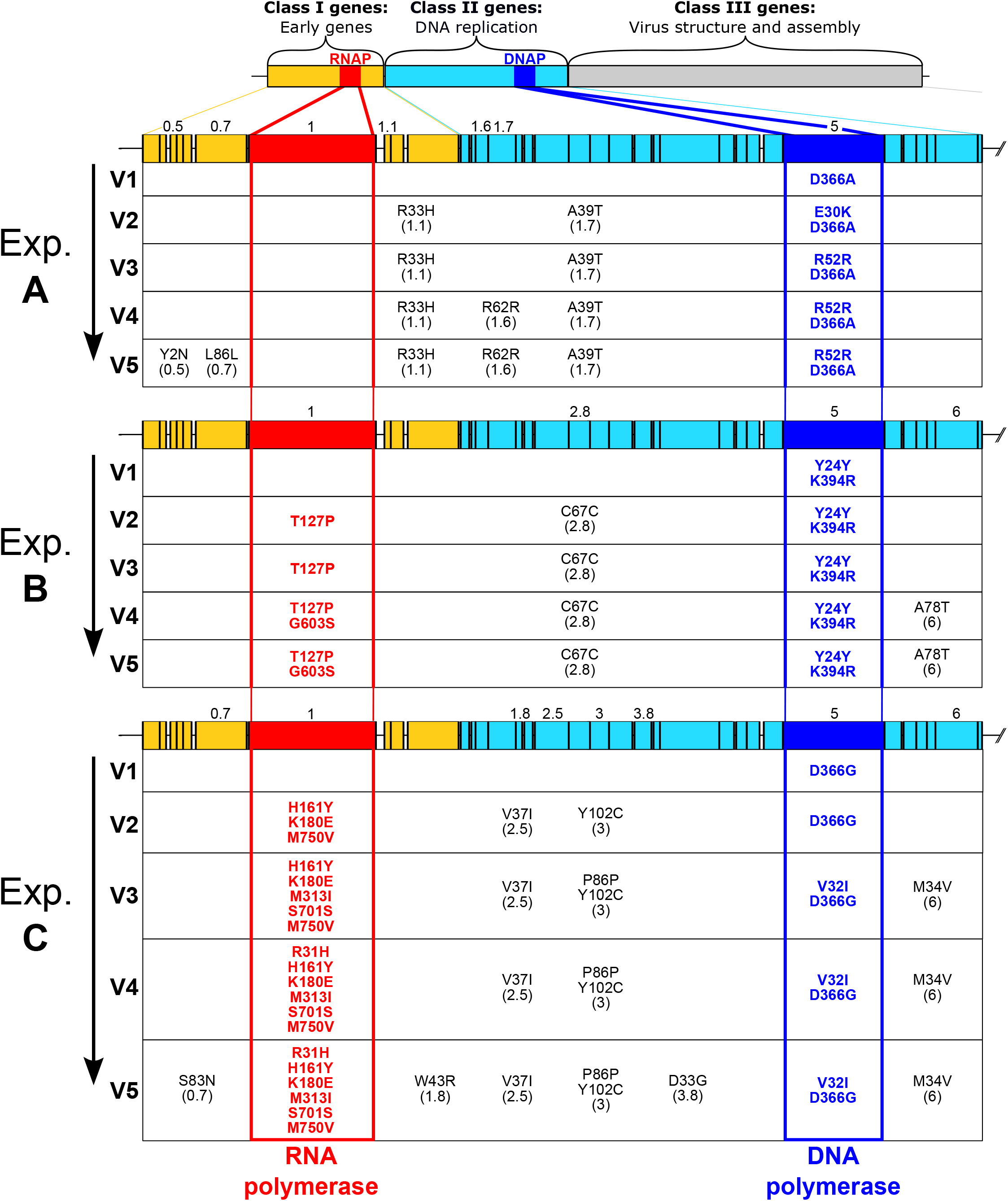
Sequence changes in the viral genome during laboratory evolution. Number of high frequency (fraction>0.95) mutations for the different viral genes during the evolution experiments of the {A,B,C} set. For the sake of clarity, mutations in class III genes are not shown here, but can be found in Figure S7. The viral DNA and RNA polymerases have been highlighted. For all other mutations, the gene is indicated within brackets. Data were obtained by Illumina sequencing of the DNA extracted after lysing of the engineered host samples (see main text and Figure 7 for details). A control experiment in which the non-evolved virus propagates in the original host yielded no mutations in the class I and class II genes and very few mutations in class III genes (see Figure S6).

**Figure 10.**
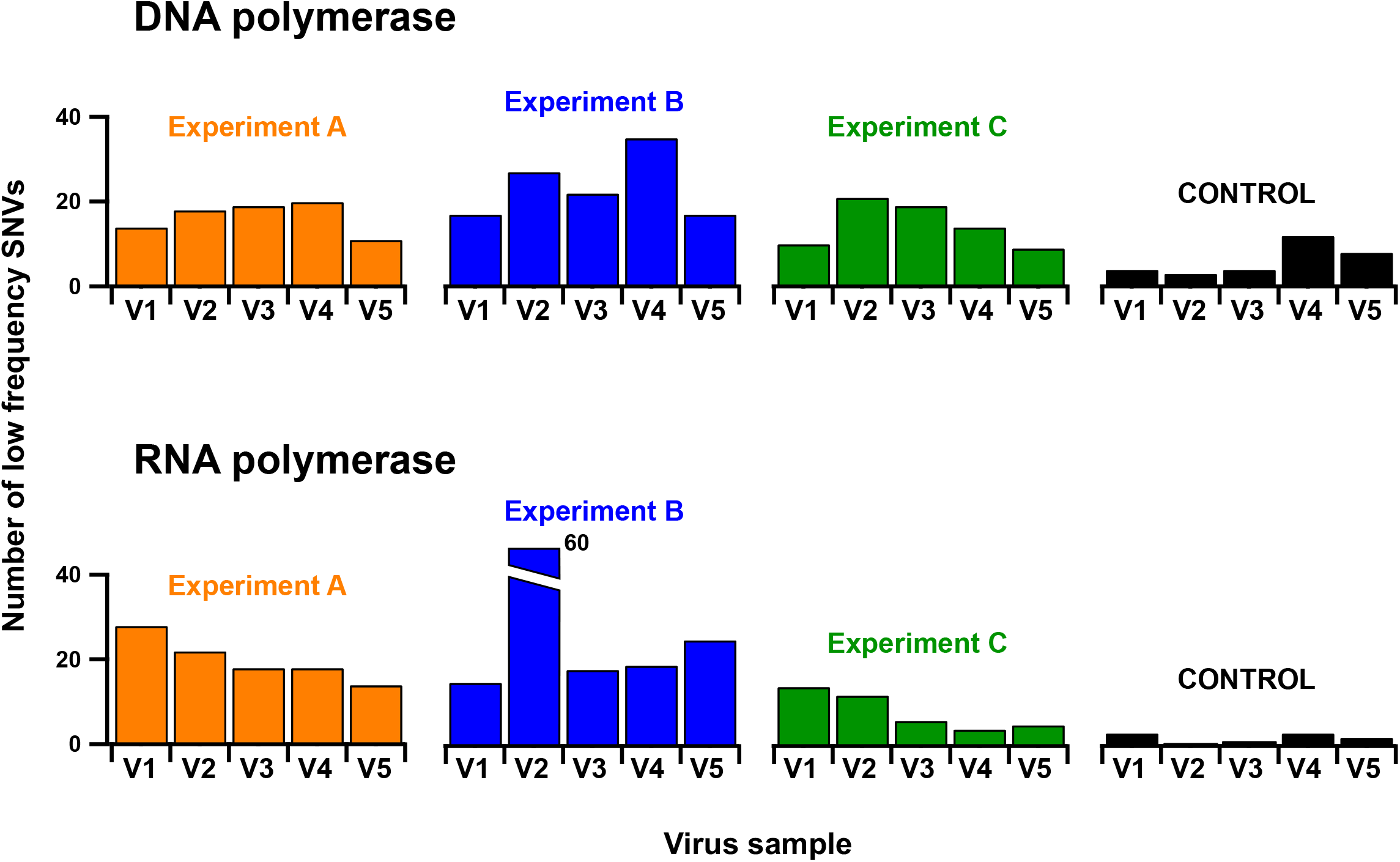
Number of low-frequency SNVs during virus adaptation to the engineered host. The numbers shown for the viral DNA and RNA polymerases were obtained from the next generation sequencing of the viral genome from the virus samples (V1-V5) at the several evolutionary rounds of the {A,B,C} set of experiments. Only total numbers of low-frequency SNV’s are shown here; see Table S2 for more detailed information. For comparison, we also include values in a control experiment in which the non-evolved phage was allowed to propagate in the original host. For the sake of clarity, the bar for the RNA polymerase from the V2 sample in experiment B has been truncated (the actual value in this case is about 60 SNV’s).

Several inferences can be made from these sequencing studies and analyses:

1. The results obtained for the DNA polymerase are consistent with those derived using Sanger sequencing discussed above. That is, the few mutations observed at high frequency do not appear in the TBD (Figure 6b) and cannot directly explain promiscuous thioredoxin recruitment. Certainly, SNVs at low frequencies occasionally occur in the TBD. However, they do so transitorily (see Table S2 for details), while any genetic mutation that directly enabled recruitment in the engineered host would have been enriched by selection. Also, no mutations are fixed in the helicase-primase (product of gene gp4) which interacts with the DNA polymerase in the replisome. Certainly, one mutation is fixed in gp2.5 during experiment C (Figure 9), but the role of this protein in the replisome is to coat the ssDNA in the lagging strand (Hamdan and Richardson, 2009). Therefore, mutations in gp2.5 cannot reasonably explain promiscuous thioredoxin recruitment by the DNA polymerase.
2. The mutations in the viral DNA polymerase increase replication errors. This is easily inferred from the numbers of low-frequency single-nucleotide variants in the virus samples from the {A,B,C} experiments as compared with a control experiment in which the non-evolved virus propagates in the control host (note that we exclude high frequency SNVs from this analysis, since their accumulation could potentially be the result of natural selection fixing mutations with a selective advantage). In fact, the pattern of much lower numbers of SNVs in the control experiments is already apparent for the DNA polymerase and RNA polymerase in Figure 10 (see also Figures S3 and S4) and it is not an artefact related to a lower number of readings in the control, as it is visually apparent in Figure S5. This result is confirmed by a more thorough analysis of the whole viral genome, as described below: We selected SNVs in the whole viral genome with frequency less than 0.2 as given by the ratio of the number of nucleotides detected as an alternative to the number of nucleotides identical to those in the reference genome in the same genomic position. To exclude variants with very low frequencies that may be due to sequencing errors, only those with a frequency greater than 0.01 were taken into account. Furthermore, to eliminate the possible noise of mutation hotspots outside the coding regions, as intergenic regions, only variants coded within the 60 genes of the virus were included. Before comparing the numbers of SNVs from different virus samples, they were normalized through division by the total number of positions read in each sample, that is, by a value that corresponds to the sample coverage (average number of reads per nucleotide) times the size of the genome. The resulting numbers of normalized alternative reads are shown in Figure 6D for the several virus samples of the {A,B,C} experiments, together with the values for a control experiment in which the non-evolved virus propagates in the original host. Lower numbers are obtained for the control as compared with any of the {A,B,C} experiments, a result which is statistically significant as supported by T Test p values of 8·10^−4^ (samples from experiment A), 10^−2^ (samples from experiment B) and 1.5·10^−3^ (samples form experiment C).
3. Virus adaptation to the engineered host correlates with mutations in the gene for the viral RNA polymerase. First, large numbers of SNVs, at both high and low frequencies, are determined for the RNA polymerase in samples from the {A,B,C} experiments, while very few SNVs are found in the control experiment using the original virus in the original host (Figures 9, 10 and S4). Again, the low number of SNVs in the control is not an artefact due to an insufficient number of readings (Figure S5). Secondly, the C>B>A engineered vs. original host adaptation pattern (Figure 8B) correlates with the number of high frequency mutations observed in the gene for viral RNA polymerase. In particular, the largest number of high-frequency mutations in the viral RNA polymerase occurs in experiment C (Figure 9), where a very efficient adaptation to the engineered host is observed (Fig. 8B). Certainly, experiment C also shows increased number of mutations in class III genes (Figures S6 and S7), but these are involved in virion assembly and host lysis and cannot be connected with replisome assembly in any reasonable way. On the other hand, as we elaborate below (see Discussion), mutations in the RNA polymerase provide a straightforward explanation for the observed virus adaptation, as they may have an effect on replisome assembly linked to the possibility of transcription errors (see Discussion for details).

### Assessing the error levels of viral RNA polymerases

We hypothesize (see Discussion for details) that transcription errors play a key role in the virus adaptation reported in this work. We deemed convenient, therefore, to qualitatively test the error level of the RNA polymerase from the last round of experiment C using a known procedure (Goldsmith and Tawfik, 2009) based on the determination of antibiotic resistance. This experiment is briefly described below:

We complemented a JM109 *E. coli* strain with two plasmids, one bearing the viral RNA polymerase gene under the promotor of the *E. coli* RNA polymerase and another one bearing the gene of TEM-1 β-lactamase under the promotor of the viral RNA polymerase gene. The viral RNA polymerase is induced by arabinose and β-lactamase is induced by IPTG. β-lactamase degrades β-lactam antibiotics and, in principle, its expression would make *E. coli* resistant to ampicillin. However, in the cells we are using for these experiments, the β-lactamase gene is transcribed by the viral RNA polymerase and, therefore, antibiotic resistance will be compromised to some extent by the error-prone nature of the RNA polymerase, as a substantial number of the phenotypic mutations caused by transcription errors will be disruptive and compromise the proper folding of β-lactamase to yield an active enzyme (Goldsmith and Tawfik, 2009). We performed experiments in which cell growth was monitored in solution upon two-fold serial dilutions of ampicillin and determined values of inhibitory ampicillin concentration (IC: maximum ampicillin concentration at which growth is observed) for both wild-type T7 RNA polymerase and the variant with the 5 non-silent mutations from the last round of experiment C (Figure 11). In the absence of arabinose induction, the viral RNA polymerase is not synthesized, the β-lactamase gene is not transcribed and, therefore, the IC value is very low and essentially identical to the value obtained using *E. coli* cells not-complemented with the plasmid bearing the β-lactamase gene. Upon arabinose (and IPTG) induction, RNA polymerase is synthesized, the β-lactamase gene is transcribed and substantial resistance to ampicillin emerges. However, the IC values are about 16-fold higher with the wild-type viral RNA polymerase as compared with the variant from evolution experiment C. This supports that the mutations fixed in the viral RNA polymerase during evolution experiment C do increase its transcription error-rate. Interestingly, Western blotting analysis (Figure 11B) suggest a somewhat higher expression level for β-lactamase when it is transcribed by RNA polymerase from the evolved virus. One possibility then is that the mutations accumulated in the viral RNA polymerase during experiment C bring about a transcription process that is both faster and more error-prone.

**Figure 11.**
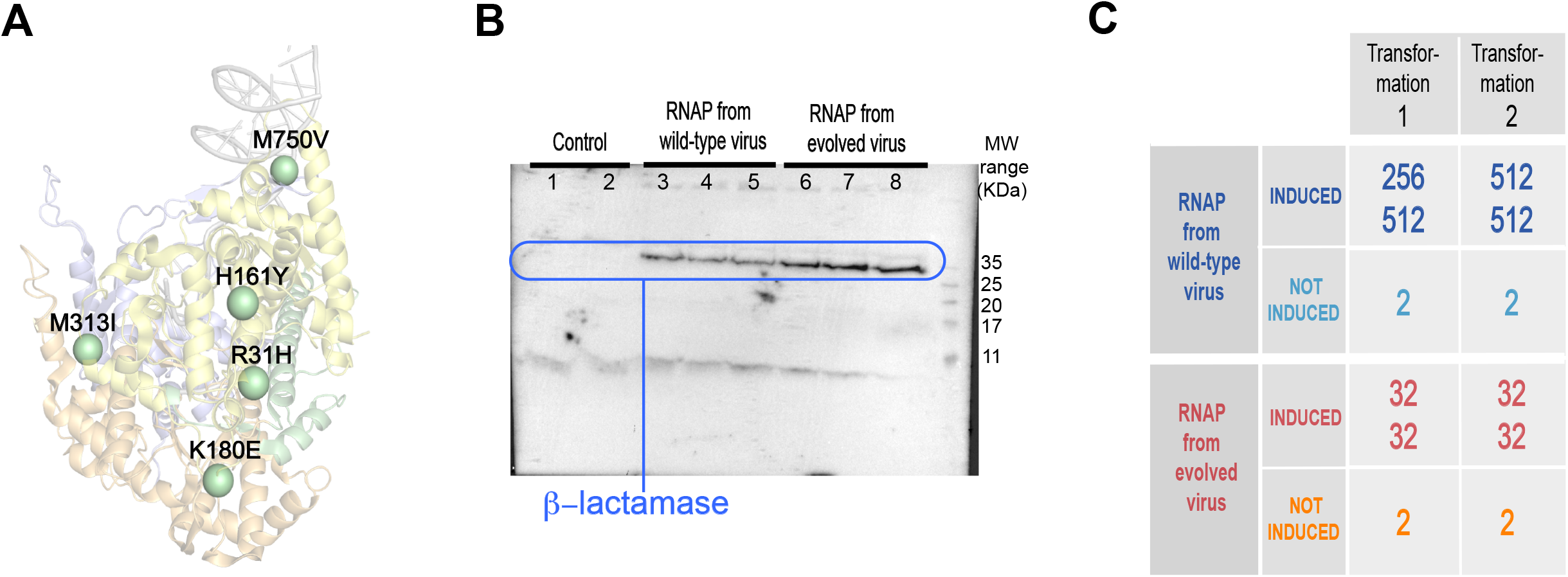
Assessment of the error level of viral RNA polymerases. **A,** 3D-structure of the viral RNA polymerase showing the positions at which non-silent mutations were fixed during evolution experiment C (see text and Figure 9 for details). **B,** Western blotting analysis of the expression of the antibiotic resistance enzyme β-lactamase when its gene is transcribed by viral RNA polymerases, both from the wild-type virus and from the evolved virus from experiment C (see text and Figure 9 for details). Three independent experiments were run with each RNA polymerase. “Control” refers to two independent experiments in which the RNA polymerase used (wild type) was not induced and, therefore the β-lactamase gene was not transcribed. **C,** Resistance to the antibiotic ampicillin of *E. coli* strains in which the gene of the antibiotic resistance enzyme β-lactamase is transcribed by viral RNA polymerases. Values of the inhibitory ampicillin concentration (IC: maximum ampicillin concentration at which growth is observed) are given for independent experiments starting with different clones from two transformations. IC values (μg/mL) are very low when the RNA polymerases are not induced (no arabinose added) and the gene of the β-lactamase is not transcribed. Upon induction of the polymerases, antibiotic resistance is observed, but lower IC values are observed with the RNA polymerase from the evolved virus (from experiment C) support an enhanced error level (see text for details). Two-fold serial dilutions of ampicillin were used in these experiments.

## DISCUSSION

### Key results from our phage evolution experiments

We challenged bacteriophage T7 to propagate in a host that had been modified to hinder the recruitment of a protein that is essential for the assembly of a functional viral replisome. The modification targets a very specific virus-host interaction involving thioredoxin (a proviral factor) and the thioredoxin binding domain (TBD) of the viral DNA polymerase. Our laboratory evolution experiments yield some surprising results. First, the virus adapted to the engineered host (i.e., it evolved to exploit the modified proviral factor) without losing its competence to propagate in the original host (i.e., it retained the capability to recruit the unmodified proviral factor). Secondly, genetic mutations cannot explain the observed adaptation. This is, of course, shown by the fact that no genetic mutations were found in the TBD domain, but it is also apparent from the observed pattern of promiscuity. Since the interaction between thioredoxin and the TBD domain of the DNA polymerase is highly specific, genetic mutations that could enable the recruitment of the modified proviral factor would likely impair the recruitment of *E. coli* thioredoxin, causing a strong trade-off, which is not observed in our evolution experiments.

### Comparison with previous reports of promiscuity in virus adaptation

It is interesting to compare our results with those recently obtained by Petrie et al. (2018) in a laboratory evolution study on the adaptation of bacteriophage λ to a new receptor. Promiscuity, (here the capability to exploit both the new and the old receptors) was also found in this study and attributed to genetic mutations that induced an ensemble of different protein conformations and generated a diversity of interaction capabilities (Petri et al., 2018). However, it is difficult to see how this mechanism could apply to our system, since no mutations at the level of DNA are fixed in the TBD domain of the viral DNA polymerase. Certainly, conformational dynamics is known to be subject to long-distance effects (Petrovic et al., 2018) and we could consider the possibility that distant mutations in the DNA polymerase caused conformational diversity in the TBD. Still, we have performed sequence determinations for a substantial number (17) of evolution experiments and have found typically 1-3 mutations per experiment, spreading overall over 21 positions in the structure of the DNA polymerase (Figure 6D). It would be unrealistic to assume that mutations at so many different positions show the same capability to trigger, through long-distance effects, the conformational heterogeneity in the TBD that it is specifically required to enable promiscuous recruitment.

### On the role of transcription errors on phage adaptation to the engineered host

Once genetic mutations in the DNA polymerase are ruled out as responsible for the observed virus adaptation, the only remaining possibility is that mutations at the phenotypic level caused by protein synthesis errors enable promiscuous thioredoxin recruitment. This possibility is reasonable, because viral replisome assembly likely needs only occur a few times per host cell to allow virus propagation and can therefore be mediated by protein variants present at very low level. Phenotypic mutations can be due to translation errors or to transcription errors. Since viruses, however, do not encode a translation machinery, the obvious suspect is the viral RNA polymerase. This enzyme (Cheetham and Steitz, 2000) transcribes the viral genes involved in virion assembly and host cell lysis (class III genes), as well as those involved in DNA replication (class II genes), including the gene for the DNA polymerase. Directed evolution studies (Brakmann and Grzeszik, 2001) indicate that, under the appropriate pressure, the T7 RNA polymerase can evolve towards increased error rates by accepting mutations in different regions of the molecule. Indeed, our experiments shown in Figure 11, in which the gene for the antibiotic resistance enzyme β-lactamase gene is transcribed by the RNA polymerase, support that the mutations fixed in the viral RNA polymerase during evolution experiment C do increase its error-rate. Overall, therefore, mutations in the gene for the viral RNA polymerase could increase transcription error rates and lead to phenotypic mutations, some of which would occur at the TBD/thioredoxin interaction region and enable recruitment. Initially, the adaptation would rely on variants of the RNA polymerase present at low level (likely the situation of experiment A), although eventually selection will lead to fixation of mutations, as it is already observed in experiments B and C (Figure 9).

### A mechanism of phage adaptation to the engineered host based on phenotypic mutations

To summarize, our results are consistent with a mechanism of adaptation to the engineered host that involves the following steps: 1) increased replication rates are brought about by mutations in the viral DNA polymerase; 2) replication errors promote genetic mutations throughout the viral genome including the gene of the viral RNA polymerase; 3) increased transcription errors cause phenotypic mutations that enable the interaction between the DNA polymerase and thioredoxin to be established in the engineered host. This mechanism provides a simple and convincing explanation for the capability of the evolved phage to propagate in both the original and the engineered hosts. Focusing for illustration on the DNA polymerase, transcription errors will generate a population of molecules with different sets of phenotypic mutations. Some of these molecules will still recruit the original thioredoxin, while some other will have the capability to recruit the alternative LPBCA thioredoxin. Note that the putative ancestral protein we have used displays higher stability (Perez-Jimenez et al., 2011) and better folding kinetics properties (Gamiz-Arco et al., 2019) than its modern *E. coli* counterpart, which should contribute to somewhat higher levels of folded protein *in vivo*. This explains that lysis times for the engineered host eventually become somewhat smaller than the lysis times for the original host in experiment C (Figure 8B).

Of course, it may seem at first surprising that, according to our proposed mechanism, phenotypic mutations enable virus replication in the engineered host while they compromise antibiotic resistance in the experiment described in Figure 11C in which the gene for the antibiotic resistance enzyme β-lactamase gene is transcribed by the RNA polymerase from the evolved virus. However, it is important to note that the required numbers of functional enzymes greatly differ between the two scenarios. Antibiotic resistance requires a substantial amount of active β-lactamase and will therefore be compromised by phenotypic mutations which, in most cases, impair proper folding to yield an active enzyme. On the other hand, virus infection of a cell typically results in the generation of a comparatively small number of virions and replication, therefore, could be achieved on the basis of just a few efficient replisomes. In this scenario, phenotypic mutations could certainly lead to misfolded proteins but also to a small, but sufficient, number of variants capable to form efficient replisomes and enable replication.

### General implications of this work

The experimental studies reported here make use of a host that has been engineered to display a very specific difference with the original host and also to promote phenotypic mutations at the targeted host-virus interaction. Still, we argue that phenotypic mutations caused by transcription errors may provide a general mechanism for virus adaptation. Virus infection of a cell typically results in the generation of a not too large number of new virions. It follows that many of the key intermolecular interactions involved in virus infection and replication need only occur a few times per host cell and could be mediated by very low levels of protein variants with enabling phenotypic mutations. To illustrate the plausible generality of the proposed adaptation mechanism, several potential examples are briefly discussed below.

Virus cross-species transmission requires that the virus has the remarkable capacity to establish key interactions in the two different molecular environments of the old and the new host. Diversity at the phenotypic level caused by transcription errors may generate an extensive capability to establish/avoid interactions in different molecular environments, thus facilitating cross-species transmission. The adaptation mechanism based on phenotypic mutations may be immediately implemented in RNA viruses in which both replication and transcription are performed by error-prone RNA-dependent RNA polymerases. For instance, RNA synthesis in coronaviruses produces, not only genomic RNA, but also subgenomic RNAs that are not encapsulated in the assembled virions, but that function as mRNAs for downstream genes (Fehr and Perlman, 2015; de Wilde et al., 2018) potentially leading to “useful” (for the virus) phenotypic diversity in other structural proteins including the spike (of which there are many copies in the virion surface). Furthermore, transcription errors may conceivably lead to viral protein variants capable of evading antiviral strategies. Phenotypic mutations might, for instance, allow the evasion of antibody neutralization, a phenomenon that plausibly contributes, for instance, to the so-called influenza puzzle (Ellebedy et al., 2016), i.e., the fact that influenza remains a health problem despite repeated exposure of the population worldwide to natural infection and to influenza viral proteins through vaccination. Another intriguing possibility is that phenotypic mutations contribute to the very low genetic barriers sometimes observed for resistance towards antivirals (Pennings, 2012; Irwin et al., 2016) by complementing the effect of the few mutations that appear at the genetic level. Obviously, these suggested possibilities rely on the notion that protein variants with suitable phenotypic mutations and present at very low concentrations may enable key processes for virus infection and replication. The interest in determining low level mutations in viral mRNAs thus emerges. This task is compromised by normal errors of next generation sequencing and the errors introduced in reverse transcription steps. Still, methods to circumvent these problems have been developed in recent years (Lu et al., 2020).

## Supporting information

Suppmental Tables and Figures

## ACKNOWLEDGEMENTS

This work was supported by Spanish Ministry of Economy and Competitiveness/FEDER Funds Grant RTI2018-097142-B-100 and by Human Frontier Science Program Grant RGP0041/2017. Viral genome library preparation and Illumina sequencing were carried out at the IPBLN Genomics Facility (CSIC, Granada, Spain) and the assistance of Dr. Alicia BarrosoDelJesus is gratefully acknowledged. We also thank Dr. John Beckwith and Dr. Dana Boyd (Harvard University) for kindly providing knockout strains used in this work.

## AUTHOR CONTRIBUTIONS

Conceptualization, B.I.-M. V.A.R., and J.M.S.-R.; Methodology, A.D., R.L.-H., V.A.R. and E.A.-L.; Experiments, R.L.-H., and V.A.R.; Formal Analyses, R.L.-H., B.I.-M., and V.A.R; Writing - Original Draft, J.M.S.-R; Writing – Review & Editing, all authors; Supervision and Funding Acquisition, J.M.S.-R.

## REFERENCES

Akabayov, B., Akabayov, S.R., Lee, S.J., Tabor, S., Kulczyk, A.W., and Richardson, C.C. (2010). Conformational dynamics of bacteriophage T7 DNA polymerase and its processivity factor, *Escherichia coli* thioredoxin. Proc. Natl. Acad. Sci. USA 107, 15033–15038.

Andrews, S. (2010). FastQC: a quality control tool for high throughput sequence data. Available online at: http://www.bioinformatics.babraham.ac.uk/projects/fastqc

Brakmann, S., and Grzeszik, S. (2001). An error-prone T7 RNA polymerase mutant generated by directed evolution. ChemBioChem 2, 212–219.

Cheetham, G.M., and Steitz, T.A. (2000). Insights into transcription: structure and function of single-subunit DNA-dependent RNA polymerases. Curr. Opin. Struct. Biol. 10, 117–123.

Cingolani, P., Platts, A., Wang, L.L., Coon, M., Nguyen, T., Wang, L., Land, S.J., Lu, X., and Ruden, D.M. (2012). A program for annotating and predicting the effects of single nucleotide polymorphisms, SnpEff: SNPs in the genome of Drosophila melanogaster strain w1118; iso-2; iso-3. Fly 6, 80–92.

Danecek, P., and McCarthy, S.A. (2017). BCFtools/csq: haplotype-aware variant consequences. Bioinformatics 33, 2037–2039.

de Wilde, A.H., Snijder, E.J., Kikkkert, M., and van Hemert, M.J. (2018). Host factors in coronavirus replication. Curr. Top. Microbiol. 419, 1–42.

Delgado, A., Arco, R., Ibarra-Molero, B., and Sanchez-Ruiz, J.M. (2017). Using resurrected ancestral proviral proteins to engineer virus resistance. Cell Reports 19, 1247–1256.

Drummond, A.D., and Wilke, C.O. (2009). The evolutionary consequences of erroneous protein synthesis. Nat. Rev. Genet. 10, 715–724.

Ellebedy, A.H., and Ahmed, R. (2016). Antiviral vaccines: challenges and advances. Chapter 15, pp 283–310, part VI of “The Vaccine Book” (second edition), Barry R. Bloom & Paul-Henri Lambert, eds. Academic Press.

Etson, C.M., Hamdan, S.M., Richardson, C.C., and van Oijen, A.M. (2010). Thioredoxin suppresses microscopic hopping of T7 DNA polymerase on duplex DNA. Proc. Natl. Acad. Sci. USA 107, 1900–1905.

Fehr, A.R., and Perlman, S. (2015). Coronaviruses: an overview of their replication and pathogenesis. Coronaviruses 1282, 1–23.

Gamiz-Arco, G., Risso, V.A., Candel, A.M., Ingles-Prieto, A., Romero-Romero, M.L., Gaucher, E.A., Gavira, J.A., Ibarra-Molero, B., and Sanchez-Ruiz, J.M. (2019). Nonconservation of folding rates in the thioredoxin family reveals degradation of ancestral unassisted-folding. Biochem. J. 476, 3631–3647.

Garrison, E., and Marth, G. (2012). Haplotype-based detection from short-read sequencing. arXiv:1207.3907.

Goldsmith, M., and Tawfik, D.S. (2009). Potential role of phenotypic mutations in the evolution of protein expression and stability. Proc. Natl. Acad. Sci. USA 106, 6197–6202.

Hamdan, S.M., and Richardson, C.C. (2009). Motors, switches and contacts in the replisome. Annu. Rev. Biochem. 78, 205–243.

Ingles-Prieto, A., Ibarra-Molero, B., Delgado-Delgado, A., Perez-Jimenez, R., Fernandez, J.M., Gaucher, E.A., Sanchez-Ruiz, E.A., and Gavira, J.A. (2013).Conservation of protein structure over four billion years. Structure 21, 1690–1697.

Irwin, K.K., Renzette, N., Kowalik, T.F., and Jensen, J.D. (2016). Antiviral drug resistance as an adaptive process. Virus Evol. 2, vew014.

Lee, S.J., and Richardson, C.C. Choreography of bacteriophage T7 DNA replication. Curr. Opin. Chem. Biol. 15, 580–586 (2011).

Li, H., and Durbin, R. (2009). Fast and accurate short read alignment with Burrows-Wheeler transform. Bioinformatics 25, 1754–60.

Lu, I.-N., Muller, C.P., and He, F.Q. (2020). Applying next-generation sequencing to unravel the mutational landscape in viral quasispecies. Virus Res. 283, 197963.

Pennings, P.S. (2012) Standing genetic variation and the evolution of drug resistance in HIV. PLoS Comput. Biol. 8, e1002527.

Perez-Jimenez R, Ingles-Prieto, A., Zao, Z.-M., Sanchez-Romero, I., Alegre-Cebollada J., Kosuri, P., Garcia-Manyes S., Kappock TJ, Tanokura, M., Holmgren A., Sanchez-Ruiz, J.M., Gaucher, E.A., and Fernandez, J.M. (2011). Single-molecule paleoenzymology probes the chemistry of resurrected enzymes. Nat. Struct. Mol. Biol. 18, 592–596.

Petrie, K.L., Palmer, N.D., Johnson, D.T., Medina, S.J., Yan, S.J., Li, V., Burmeister, A.R., and Meyer, J.R. (2018). Destabilizing mutations encode nongenetic variation that drives evolutionary innovation. Science 359, 1542–1545.

Petrovic, D., Risso, V.A., Kamerlin, S.C.L., and Sanchez-Ruiz, J.M. (2018). Conformational dynamics and enzyme evolution. J. R. Soc. Interface 15, 20180330.

Seemann, T. (2015). Snippy-Rapid haploid variant calling and core SNP phylogeny. GitHub. Available at: https://github.com/tseemann/snippy/.

Traverse, C.C., and Ochman, H. (2016). Conserved rates and patterns of transcription errors across bacterial growth states and lifestyles. Proc. Natl. Acad. Sci. USA 113, 3311–3316.

Whitehead, D.J., Wilke, C.O., Vernazobres, D., and Bornberg-Bauer, E. (2008). The look-ahead effect of phenotypic mutations. Biol. Direct 3, 18.

