## Supplementary material for "Evidence for a role of phenotypic mutations in virus adaptation": Suppmental Tables and Figures

*^1^Departamento de Quimica Fisica. Facultad de Ciencias, Unidad de Excelencia de Quimica Aplicada a Biomedicina y Medioambiente (UEQ), Universidad de Granada, 18071 Granada, Spain.*

*^2^Unidad de Bioinformática. Instituto de Parasitología y Biomedicina “López Neyra”, CSIC, 18016 Armilla, Granada, Spain.*

^†^These authors contributed equally to this work

**METHODS**

**Strains used in this work**

The *E. coli* strains used in this work have been previously described in detail (Delgado et al., 2017). Briefly, we use as the original host the *E. coli* strain DHB4. The engineered host is based upon a receptor strain FA41 which is a DHB4 strain deficient in the two thioredoxins identified in the cytoplasm of *E. coli*: thioredoxin 1, which we refer throughout the text simply as *E. coli* thioredoxin and which is the thioredoxin recruited by the non-evolved virus for its replisome, and thioredoxin 2, which is induced under some stress conditions and has an additional zinc-binding domain. This DHB4 *trxA trxC* strain was a gift from Jon Beckwith (Harvard Medical School). As previously described (Delgado et al., 2017), we further performed the curation of the F’ factor (which would prevent phage infection) and lysogenization (using the λDE3 Lysogenization kit from Novagen) to introduce in the bacterial chromosome the gene of the T7 RNA polymerase. This results in the *E. coli* Trx^–^ strain that has been used as a control throughout this work. The gene for LPBCA thioredoxin was cloned in pET30a(+) under a T7 RNA polymerase promoter as previously described (Delgado et al.). This system is leaky and leads to a basal expression level, even under non-inducing conditions. Complementation of the *E. coli* Trx^–^ strain with this plasmid produced the engineered host extensively studied in this work. The gene for *E. coli* thioredoxin was also cloned in pET30a(+) and complementation of the *E. coli* Trx^–^ strain with this plasmid produced the version of the original host used in this work.

**Plaque assays**

Plaque assays were performed as described previously in detail (Delgado et al., 2017). Briefly, phage samples were serially diluted in Tris buffer 20 mM pH 7.4 (including 100 mM NaCl, 10 mM MgSO_4_). Strains were grown to an absorbance of ~0.5 in LB medium at 37 ºC and 100 μL of the cell suspension were mixed with 100 μL of a phage suspension and the resulting mix was combined with 4 mL of molten agar. Plates were incubated at 37 ºC overnight or 15 hours for the experiments A, B and C (Fig. 3). Numbers of plaques were determined from visual inspection and counting. Phage titer was calculated upon serial dilution experiments from the number of plaques at the two highest phage dilutions at which plaques are observed.

**Lysis experiments and DNA extraction**

Lysis experiments were carried out using a straightforward modification of the protocol we previously used for the determination of generation times (Delgado et al., 2017). Briefly, preinocula in LB were incubated overnight at 37 ºC. Cultures were then diluted 1/200 in fresh medium and the absorbance at 600 nm was determined as function of time for about 4 hours typically while keeping the temperature at 37 ºC. A volume of virus sample was added when the absorbance was 0.25-0.3. After lysis, cultures were collected and kept at -20 ºC until DNA extraction. For this, samples were thawed on ice, mixed with 1/60 chloroform, vortexed and centrifuged at 4000g for 10 minutes. DNA was isolated from the supernatant using the phage DNA isolation kit from Norgen (CAT#46800).

**Evolution experiments**

The evolution experiments were initiated by a standard plaque assay in which the host engineered strain was infected with an appropriate dilution of wild type phage suspension. Initially, after overnight incubation a clearly well-isolated large plaque was picked up from the agar plate and transferred to a 1 mL of buffer (Tris buffer 20 mM pH 7.4, 100 mM NaCl, 10 mM MgSO_4_). Virus particles were allowed to diffuse for 2 hours and titration of the phage suspension was performed. Subsequently, one 100 μl aliquot was used to start the next round of evolution by infection of a fresh culture of the engineered host. An additional amplification step (Delgado et al., 2017) was included in the rounds before the one-month break in the experiment shown in the lower panel of Figure 4.

**Genomic DNA sequencing**

gDNA samples from ~50 isolates of bacteriophage T7 were extracted with the phage isolation kit from Norgen (CAT#46800) and quantified using a Nanodrop One (ThermoFisher). Subsequently, gDNA was purified using MucleoMag Beads (Omega) without size selection. The quality of the gDNA was evaluated by Qubit dsDNA HS Assay kit (ThermoFisher) and 0.8% agarose gel electrophoresis. 37 libraries were constructed with an input of 100-300 ng of DNA using the NexteraTM DNA Flex Library Preparation kit with only 5 PCR cycles in the indexing step. The quality of the libraries was validated by the Qubit dsDNA HS Assay kit (ThermoFisher) with a 2100 Bioanalyzer (Agilent Technologies). Libraries were sequenced on an Illumina MiSeq producing 1170569 of 2x250 bp read, i.e., 61762 average reads per sample, reaching an average coverage of 369x.

Initially, the sequences for each isolate were subjected to quality evaluation using FASTQC software, which provides an extensive report of the quality of the reads (Andrews, 2010). On the basis of the generated report, we determined a very high drip in the quality value of the last sequenced nucleotide, which was, therefore, eliminated. We also verified that the amount of Ns and adapters included in the sequences was almost negligible. The average quality of all reads was above 30. After this quality filter, we called and annotated all variants on the basis of the following steps performed by Snippy (Seemann, 2015). Burrow Wheeler Alignment (Li and Durbin, 2009) was used to align the reads against the reference genome (GCF_000844825 from GeneBank). Samtools (Li and Durbin, 2009) was used to sort, mark and remove duplicate sequences. Variant calling was performed by Freebayes (Garrison and Marth, 2012) using the following parameters: “-P 0 – C 10 –min-repeat-entropy 1.5 –strict-vcf –q 15 –m 30” which involves the exclusion of reads having a mapping quality of less than 30. Furthermore, alleles were excluded from the analysis if their supporting base quality was less than 15. The remaining variants were processed by snpEff (Cingolani et al., 2012) to annotate and predict their effects on genes and proteins. The single nucleotide variants discussed in the text are the outcome of an additional filtering process by bcftools (Danecek and McCarthy, 2017) to eliminate further possible errors. Variant determination at a given position required at least 10 sequences including the position and a mapping quality of at least 100. Finally, we eliminated all those variants that did not have at least 2 reads supporting the alternative nucleotide or that had a frequency lower than 0.01 (the frequency is calculated by dividing the number of reads that support the alternative by the number of reads whole nucleotide is equal to the reference). Raw sequencing data are available in the Sequence Read Archive (SRA) under the PRJNA656432 BioProject accession number.

**Antibiotic-resistance assessment of RNA polymerase error level**

We used *E. coli* JM109 competent cells (Agilent), the following plasmids,

pACYC -T7 RNAP-wt, encoding the wild type RNA polymerase under lacI repression and induced by addition of L-arabinose,

pACYC -T7 RNAP-variant, encoding the RNA polymerase from the viral V5 sample of experiment C, under lacI repression and induced by addition of L-arabinose,

pET30a(+)-TEM-1, encoding the TEM-1 β-lactamase gene under the control of the T7 operon and induced by addition of IPTG,

and the following media for bacterial growth,

LB medium. 10 g/L Bacto -Tryptone, 5 g/L Bacto -Yeast Extract, 5 g/L NaCl, autoclave to sterilize. Allow the auto-claved medium to cool to 55ºC and add chloramphenicol (35 mg/mL), kanamycine (50 mg/mL) and glucose (1%). For LB plates, 1.5% Bacto-agar (15 g/L) was added prior to autoclaving.

SOC medium. 2% Tryptone (bacto), 0.5% Yeast Extract, 10 mM NaCl, 2.5 mM MgCl_2_, 10 mM MgSO_4_, 20 mM glucose.

Transformation of plasmid DNA into *E. coli* using the heat shock method was performed using well-established protocols. Briefly, after a short incubation in ice (30 min), a mixture of chemically competent bacteria and DNA is placed at 42°C for 45 seconds (heat shock) and then placed back in ice. SOC media is added and the transformed cells are incubated at 37°C for 1 h with agitation. Cells were plated on LB agar plates with chloramphenicol (35 μg/mL), kanamycine (50 μg/mL) and glucose (1%) and grown without selection (O/N, 37 °C). Colonies carrying a TEM-1 and T7 RNAP-wt or TEM-1 and T7 RNAP-variant were picked and individually grown in 5 mL of “Liquid growth media” [LB medium with chloramphenicol (35 μg/mL), kanamycine (50 μg/mL) and glucose (0.2%)] for 16 h at 37 °C with shaking.

To test ampicillin resistance, cultures were diluted 1/100 in fresh ‘‘Liquid growth media” and strains were grown to an absorbance of ~0.2 add L-arabinose to be concentration final 0.2%, and culture at 37ºC for 5 h. Then, β-lactamase expression was induced by addition of 0.5 mM IPTG for 2 h (37 °C, shaking). Finally, after de induction the cultures were diluted (1:100) in fresh “Liquid growth media” with IPTG 0.5 mM and varying concentrations of ampicillin (0, 2, 4, 8, 16, 32, 64,128, 256, 512, 1024, 2048 μg/mL). Culture were grown overnight at 37 °C and the inhibitory concentration for TEM-1 were determined from visual inspection. The inhibitory concentration of each transformant, were assayed in duplicate with two independent transformations.

To assess the expression levels of TEM-1 β-lactamase when its gene is transcribed by the viral RNA polymerases, cultures (prepared as described above) were diluted 1/100 in fresh ‘‘Liquid growth media”, the strains were grown to an absorbance of ~0.2, L-arabinose to be concentration final 0.2% was added and the cultures were grown at 37ºC for 5 h. β-lactamase expression was then induced by addition of 0.5 mM IPTG. After 2 hours at 37 °C with shaking, the cultures were diluted (1:100) in fresh “Liquid growth media” with IPTG 0.5 mM and grown overnight at 37 °C. 5 ml of bacterial cultures were then centrifuged for 10 min at 4000 rpm at 4ºC. Pellets were resuspended in 2 ml buffer [10 mM potassium phosphate buffer, pH 6.5, 1 mM EDTA, 20% (w/v) sucrose, 1 mg ml^-1^ lysozyme]. Cell lysis was performed by sonication using a Vibra cell ultrasonics processor (1 seg. pulse, 60% amplitude, eight cycles) with cooling. Equal volumes of cell lysis preps were loaded onto a 15% PAGE-gel. The cell lysis preparations were analysed by Western blotting. Western blots were revealed with monoclonal Anti-TEM-1 β-lactamase antibody produced in mouse (Invitrogen MA1-10712) and Horseradish Peroxidase (HRP)-conjugated goat anti-mouse antibodies (Santa Cruz Biotechnology ). After chemiluminescence (ECL) detection, the membrane was photographed.

**Table S1. Mutations in the viral DNA polymerase determined from PCR and Sanger sequencing on samples from the early rounds of 14 evolution experiments**

| Experiment | POS | NT_CHANGE | AA_CHANGE | V1 | V2 | V3 |
| --- | --- | --- | --- | --- | --- | --- |
| 1 | 15533 | 1181A>G | Lys394Arg | X | X | ND |
| 2 | 14473 | 121G>T | Ala41Ser | X | ND | X |
| 2 | 15652 | 1300T>C | Phe434Leu | - | ND | X |
| 3 | 14398 | 46A>G | Thr16Ala | X | ND | X |
| 4 | 14485 | 133G>A | Ala45Thr | X | ND | X |
| 5 | 14463 | 111T>C | Ser37Ser | X | ND | X |
| 6 | 14382 | 30C>T | Ala10Ala | X | ND | X |
| 6 | 15445 | 1093G>A | Asp365Asn | - | ND | X |
| 6 | 15578 | 1226A>G | Tyr409Cys | - | ND | X |
| 7 | 14434 | 82A>G | Thr28Ala | X | ND | X |
| 8 | 14495 | 143C>T | Ala48Val | X | X | ND |
| 8 | 15512 | 1160A>G | Lys387Arg | X | X | ND |
| 9 | 14485 | 133G>A | Ala45Thr | X | X | ND |
| 9 | 15512 | 1160A>G | Lys387Arg | X | X | ND |
| 10 | 14434 | 82A>G | Thr28Ala | X | - | ND |
| 11 | 14359 | 7G>A | Val3Ile | X | X | ND |
| 11 | 15533 | 1181A>G | Lys394Arg | X | X | ND |
| 12 | 14412 | 60C>T | Cys20Cys | X | X | ND |
| 12 | 15380 | 1028A>G | Gln343Arg | X | X | ND |
| 13 | 14494 | 142G>A | Ala48Val | X | X | ND |
| 13 | 15512 | 1160A>G | Lys387Arg | X | X | ND |
| 14 | 14463 | 111T>C | Ser37Ser | X | X | ND |
| 14 | 15449 | 1097A>C | Asp366Ala | X | X | ND |

**Table S2. Single nucleotide variants determined for viral DNA and RNA polymerases. Illumina sequencing was performed on DNA extracted from virus samples from the evolution the {A,B,C} set of evolution experiments. The numbers of reading for the alternative and the reference nucleotide are given. Mutations that became fixated are highlighted in yellow. SNVs in the TBD domain of the RNA polymerase are shown in bold.**

**EXPERIMENT A**

| **POS** | **NT_CHANGE** | **AA_CHANGE** | **LOCUS_TAG** | **GENE** | **PRODUCT** | **V1** | **V2** | **V3** | **V4** | **V5** |
| --- | --- | --- | --- | --- | --- | --- | --- | --- | --- | --- |
| 3187 | 17T>G | Ile6Ser | T7p07 | gene 1 | T3/T7-like RNA polymerase |  |  |  |  | 2/155 |
| 3208 | 38A>C | Asp13Ala | T7p07 | gene 1 | T3/T7-like RNA polymerase |  |  |  |  | 2/153 |
| 3233 | 63C>T | Phe21Phe | T7p07 | gene 1 | T3/T7-like RNA polymerase |  |  | 2/178 |  |  |
| 3246 | 76G>A | Asp26Asn | T7p07 | gene 1 | T3/T7-like RNA polymerase |  |  | 2/183 |  |  |
| 3249 | 79C>T | His27Tyr | T7p07 | gene 1 | T3/T7-like RNA polymerase |  | 2/175 |  |  |  |
| 3271 | 101G>A | Arg34His | T7p07 | gene 1 | T3/T7-like RNA polymerase |  | 2/163 |  |  |  |
| 3291 | 121C>T | His41Tyr | T7p07 | gene 1 | T3/T7-like RNA polymerase |  |  | 2/166 |  |  |
| 3301 | 131A>G | Tyr44Cys | T7p07 | gene 1 | T3/T7-like RNA polymerase |  |  | 5/171 |  |  |
| 3311 | 141T>C | Gly47Gly | T7p07 | gene 1 | T3/T7-like RNA polymerase |  | 2/174 |  |  |  |
| 3326 | 156C>A | Arg52Arg | T7p07 | gene 1 | T3/T7-like RNA polymerase | 2/146 |  |  |  |  |
| 3343 | 173A>G | Gln58Arg | T7p07 | gene 1 | T3/T7-like RNA polymerase |  |  | 2/175 |  |  |
| 3403 | 233T>G | Leu78Arg | T7p07 | gene 1 | T3/T7-like RNA polymerase | 2/138 |  |  |  |  |
| 3429 | 259G>A | Asp87Asn | T7p07 | gene 1 | T3/T7-like RNA polymerase | 2/153 |  |  |  |  |
| 3441 | 271G>T | Glu91* | T7p07 | gene 1 | T3/T7-like RNA polymerase |  |  | 2/161 |  |  |
| 3442 | 272A>C | Glu91Ala | T7p07 | gene 1 | T3/T7-like RNA polymerase | 2/155 |  |  |  |  |
| 3482 | 312G>A | Gln104Gln | T7p07 | gene 1 | T3/T7-like RNA polymerase | 2/151 |  |  |  |  |
| 3508 | 338C>T | Ala113Val | T7p07 | gene 1 | T3/T7-like RNA polymerase |  | 30/157 |  |  |  |
| 3510 | 340G>A | Val114Ile | T7p07 | gene 1 | T3/T7-like RNA polymerase |  |  | 2/148 |  |  |
| 3532 | 362C>T | Thr121Ile | T7p07 | gene 1 | T3/T7-like RNA polymerase | 2/155 |  |  |  |  |
| 3550 | 380C>T | Thr127Ile | T7p07 | gene 1 | T3/T7-like RNA polymerase |  | 2/162 |  |  |  |
| 3560 | 390C>A | Asp130Glu | T7p07 | gene 1 | T3/T7-like RNA polymerase | 2/153 |  |  |  |  |
| 3588 | 418G>A | Ala140Thr | T7p07 | gene 1 | T3/T7-like RNA polymerase | 2/146 |  |  |  |  |
| 3598 | 428G>A | Arg143Gln | T7p07 | gene 1 | T3/T7-like RNA polymerase | 2/147 |  |  |  |  |
| 3636 | 466G>T | Asp156Tyr | T7p07 | gene 1 | T3/T7-like RNA polymerase | 4/124 |  |  |  |  |
| 3636 | 466G>A | Asp156Asn | T7p07 | gene 1 | T3/T7-like RNA polymerase |  |  |  |  | 2/120 |
| 3640 | 470T>C | Leu157Pro | T7p07 | gene 1 | T3/T7-like RNA polymerase |  |  |  |  | 2/129 |
| 3793 | 623A>T | Asp208Val | T7p07 | gene 1 | T3/T7-like RNA polymerase |  |  | 2/153 |  |  |
| 3849 | 679G>T | Val227Phe | T7p07 | gene 1 | T3/T7-like RNA polymerase | 2/104 |  |  |  |  |
| 3876 | 706G>A | Val236Ile | T7p07 | gene 1 | T3/T7-like RNA polymerase |  | 2/172 |  |  |  |
| 3884 | 714T>C | Gly238Gly | T7p07 | gene 1 | T3/T7-like RNA polymerase |  |  |  | 2/138 |  |
| 3887 | 717A>G | Gln239Gln | T7p07 | gene 1 | T3/T7-like RNA polymerase |  |  |  | 2/139 |  |
| 3903 | 733G>A | Glu245Lys | T7p07 | gene 1 | T3/T7-like RNA polymerase |  |  |  | 2/141 |  |
| 3912 | 742C>T | Pro248Ser | T7p07 | gene 1 | T3/T7-like RNA polymerase | 2/117 |  |  |  |  |
| 3917 | 747A>G | Glu249Glu | T7p07 | gene 1 | T3/T7-like RNA polymerase |  |  | 2/175 |  |  |
| 3925 | 755A>G | Glu252Gly | T7p07 | gene 1 | T3/T7-like RNA polymerase |  |  | 2/171 |  |  |
| 3950 | 780G>A | Ala260Ala | T7p07 | gene 1 | T3/T7-like RNA polymerase |  | 2/163 |  |  |  |
| 3988 | 818T>C | Val273Ala | T7p07 | gene 1 | T3/T7-like RNA polymerase |  | 2/156 |  |  |  |
| 4194 | 1024A>C | Thr342Pro | T7p07 | gene 1 | T3/T7-like RNA polymerase |  |  |  | 14/129 |  |
| 4222 | 1052A>G | Asp351Gly | T7p07 | gene 1 | T3/T7-like RNA polymerase | 2/192 |  |  |  |  |
| 4262 | 1092G>A | Pro364Pro | T7p07 | gene 1 | T3/T7-like RNA polymerase |  |  | 7/160 |  |  |
| 4289 | 1119T>C | Ala373Ala | T7p07 | gene 1 | T3/T7-like RNA polymerase |  |  | 2/159 |  |  |
| 4310 | 1140T>C | Ala380Ala | T7p07 | gene 1 | T3/T7-like RNA polymerase |  | 2/197 |  |  |  |
| 4314 | 1144G>A | Ala382Thr | T7p07 | gene 1 | T3/T7-like RNA polymerase |  |  | 2/153 |  |  |
| 4333 | 1163A>T | Asp388Val | T7p07 | gene 1 | T3/T7-like RNA polymerase | 2/176 |  |  |  |  |
| 4346 | 1176G>A | Lys392Lys | T7p07 | gene 1 | T3/T7-like RNA polymerase |  |  |  | 2/117 |  |
| 4393 | 1223T>A | Phe408Tyr | T7p07 | gene 1 | T3/T7-like RNA polymerase |  | 2/158 |  |  |  |
| 4398 | 1228A>G | Asn410Asp | T7p07 | gene 1 | T3/T7-like RNA polymerase |  | 3/160 |  |  |  |
| 4428 | 1258A>G | Met420Val | T7p07 | gene 1 | T3/T7-like RNA polymerase | 2/173 |  |  |  |  |
| 4499 | 1329G>T | Leu443Leu | T7p07 | gene 1 | T3/T7-like RNA polymerase |  | 2/146 |  |  |  |
| 4501 | 1331T>C | Leu444Pro | T7p07 | gene 1 | T3/T7-like RNA polymerase |  |  | 2/130 |  |  |
| 4535 | 1365A>T | Glu455Asp | T7p07 | gene 1 | T3/T7-like RNA polymerase |  | 2/152 |  |  |  |
| 4541 | 1371C>T | Tyr457Tyr | T7p07 | gene 1 | T3/T7-like RNA polymerase |  | 2/161 |  |  |  |
| 4605 | 1435A>T | Ile479Phe | T7p07 | gene 1 | T3/T7-like RNA polymerase |  |  |  |  | 2/117 |
| 4618 | 1448A>G | Glu483Gly | T7p07 | gene 1 | T3/T7-like RNA polymerase |  |  | 2/141 |  |  |
| 4619 | 1449G>A | Glu483Glu | T7p07 | gene 1 | T3/T7-like RNA polymerase | 2/181 |  |  |  |  |
| 4641 | 1471G>A | Ala491Thr | T7p07 | gene 1 | T3/T7-like RNA polymerase |  |  |  |  | 20/111 |
| 4644 | 1474T>C | Cys492Arg | T7p07 | gene 1 | T3/T7-like RNA polymerase |  |  |  |  | 2/113 |
| 4645 | 1475G>A | Cys492Tyr | T7p07 | gene 1 | T3/T7-like RNA polymerase |  |  |  | 2/104 |  |
| 4668 | 1498A>T | Thr500Ser | T7p07 | gene 1 | T3/T7-like RNA polymerase | 2/178 |  |  |  |  |
| 4671 | 1501T>A | Trp501Arg | T7p07 | gene 1 | T3/T7-like RNA polymerase | 2/174 |  |  |  |  |
| 4695 | 1525T>A | Phe509Ile | T7p07 | gene 1 | T3/T7-like RNA polymerase |  |  |  | 2/102 |  |
| 4696 | 1526T>C | Phe509Ser | T7p07 | gene 1 | T3/T7-like RNA polymerase | 2/166 |  |  |  |  |
| 4700 | 1530C>A | Cys510* | T7p07 | gene 1 | T3/T7-like RNA polymerase |  |  |  | 2/101 |  |
| 4729 | 1559G>A | Gly520Glu | T7p07 | gene 1 | T3/T7-like RNA polymerase |  |  |  | 2/109 |  |
| 4741 | 1571A>G | His524Arg | T7p07 | gene 1 | T3/T7-like RNA polymerase |  |  |  | 2/108 |  |
| 4742 | 1572C>T | His524His | T7p07 | gene 1 | T3/T7-like RNA polymerase |  |  |  | 2/108 |  |
| 4749 | 1579A>G | Ser527Gly | T7p07 | gene 1 | T3/T7-like RNA polymerase |  | 2/146 |  |  |  |
| 4800 | 1630C>T | Gln544* | T7p07 | gene 1 | T3/T7-like RNA polymerase |  |  |  | 2/110 |  |
| 4801 | 1631A>G | Gln544Arg | T7p07 | gene 1 | T3/T7-like RNA polymerase |  |  |  |  | 2/115 |
| 4802 | 1632G>C | Gln544His | T7p07 | gene 1 | T3/T7-like RNA polymerase | 2/185 |  |  |  |  |
| 4853 | 1683G>A | Leu561Leu | T7p07 | gene 1 | T3/T7-like RNA polymerase |  | 2/161 |  |  |  |
| 4893 | 1723G>A | Ala575Thr | T7p07 | gene 1 | T3/T7-like RNA polymerase |  |  |  |  | 2/130 |
| 4976 | 1806T>C | Thr602Thr | T7p07 | gene 1 | T3/T7-like RNA polymerase |  | 2/178 |  |  |  |
| 4996 | 1826T>A | Val609Asp | T7p07 | gene 1 | T3/T7-like RNA polymerase |  |  |  |  | 2/121 |
| 5064 | 1894C>A | Arg632Ser | T7p07 | gene 1 | T3/T7-like RNA polymerase | 2/174 |  |  |  |  |
| 5087 | 1917C>T | Tyr639Tyr | T7p07 | gene 1 | T3/T7-like RNA polymerase | 2/159 |  |  |  |  |
| 5170 | 2000T>C | Phe667Ser | T7p07 | gene 1 | T3/T7-like RNA polymerase |  |  |  |  | 2/120 |
| 5173 | 2003C>A | Thr668Asn | T7p07 | gene 1 | T3/T7-like RNA polymerase | 2/162 |  |  |  |  |
| 5212 | 2042T>C | Ile681Thr | T7p07 | gene 1 | T3/T7-like RNA polymerase |  |  |  | 2/146 |  |
| 5216 | 2046G>A | Trp682* | T7p07 | gene 1 | T3/T7-like RNA polymerase |  | 2/142 |  |  |  |
| 5239 | 2069T>C | Val690Ala | T7p07 | gene 1 | T3/T7-like RNA polymerase |  |  |  | 2/154 |  |
| 5251 | 2081A>G | Glu694Gly | T7p07 | gene 1 | T3/T7-like RNA polymerase |  |  |  |  | 2/119 |
| 5257 | 2087T>C | Met696Thr | T7p07 | gene 1 | T3/T7-like RNA polymerase |  |  |  | 2/147 |  |
| 5288 | 2118G>A | Leu706Leu | T7p07 | gene 1 | T3/T7-like RNA polymerase |  |  |  | 2/138 |  |
| 5340 | 2170G>A | Ala724Thr | T7p07 | gene 1 | T3/T7-like RNA polymerase | 2/149 |  |  |  |  |
| 5374 | 2204T>A | Val735Glu | T7p07 | gene 1 | T3/T7-like RNA polymerase |  |  |  | 2/132 |  |
| 5376 | 2206T>C | Trp736Arg | T7p07 | gene 1 | T3/T7-like RNA polymerase |  |  |  | 2/134 |  |
| 5379 | 2209C>T | Gln737* | T7p07 | gene 1 | T3/T7-like RNA polymerase |  |  | 2/146 |  |  |
| 5389 | 2219A>C | Lys740Thr | T7p07 | gene 1 | T3/T7-like RNA polymerase |  | 2/152 |  |  |  |
| 5413 | 2243A>G | Asn748Ser | T7p07 | gene 1 | T3/T7-like RNA polymerase |  |  |  |  | 2/104 |
| 5427 | 2257G>A | Gly753Ser | T7p07 | gene 1 | T3/T7-like RNA polymerase |  |  | 2/145 |  |  |
| 5439 | 2269T>C | Leu757Leu | T7p07 | gene 1 | T3/T7-like RNA polymerase | 2/150 |  |  |  |  |
| 5499 | 2329G>A | Gly777Ser | T7p07 | gene 1 | T3/T7-like RNA polymerase |  |  | 2/147 |  |  |
| 5501 | 2331T>C | Gly777Gly | T7p07 | gene 1 | T3/T7-like RNA polymerase |  | 2/160 |  |  |  |
| 5515 | 2345T>C | Phe782Ser | T7p07 | gene 1 | T3/T7-like RNA polymerase | 2/159 |  |  |  |  |
| 5574 | 2404T>C | Tyr802His | T7p07 | gene 1 | T3/T7-like RNA polymerase |  | 2/150 |  |  |  |
| 5605 | 2435A>T | Asp812Val | T7p07 | gene 1 | T3/T7-like RNA polymerase |  |  |  | 2/133 |  |
| 5642 | 2472G>A | Leu824Leu | T7p07 | gene 1 | T3/T7-like RNA polymerase | 2/162 |  |  |  |  |
| 5643 | 2473T>C | Phe825Leu | T7p07 | gene 1 | T3/T7-like RNA polymerase |  | 2/150 |  |  |  |
| 5644 | 2474T>C | Phe825Ser | T7p07 | gene 1 | T3/T7-like RNA polymerase | 2/163 |  |  |  |  |
| 5675 | 2505A>G | Thr835Thr | T7p07 | gene 1 | T3/T7-like RNA polymerase |  |  | 6/124 |  |  |
| 5723 | 2553C>T | Asp851Asp | T7p07 | gene 1 | T3/T7-like RNA polymerase |  |  |  |  | 2/121 |
| 5739 | 2569C>T | Gln857* | T7p07 | gene 1 | T3/T7-like RNA polymerase | 2/161 |  |  |  |  |
| 5755 | 2585C>T | Pro862Leu | T7p07 | gene 1 | T3/T7-like RNA polymerase |  |  | 2/134 |  |  |
| 5770 | 2600A>G | Lys867Arg | T7p07 | gene 1 | T3/T7-like RNA polymerase |  | 2/170 |  |  |  |
| 5799 | 2629G>C | Glu877Gln | T7p07 | gene 1 | T3/T7-like RNA polymerase |  |  |  | 2/144 |  |
| 14367 | 15C>T | Asp5Asp | T7p29 | gene 5 | DNA polymerase |  |  |  | 2/152 |  |
| 14374 | 22G>A | Ala8Thr | T7p29 | gene 5 | DNA polymerase |  |  | 2/146 |  |  |
| 14394 | 42C>A | Ser14Arg | T7p29 | gene 5 | DNA polymerase | 2/180 |  |  |  |  |
| 14417 | 65T>C | Val22Ala | T7p29 | gene 5 | DNA polymerase |  | 2/169 |  |  |  |
| 14425 | 73G>A | Asp25Asn | T7p29 | gene 5 | DNA polymerase |  |  | 3/164 |  |  |
| 14428 | 76T>C | Tyr26His | T7p29 | gene 5 | DNA polymerase |  | 2/177 |  |  |  |
| 14434 | 82A>G | Thr28Ala | T7p29 | gene 5 | DNA polymerase |  |  |  |  | 2/99 |
| 14440 | 88G>A | Glu30Lys | T7p29 | gene 5 | DNA polymerase |  | 161/169 | 2/163 |  |  |
| 14487 | 135G>A | Ala45Ala | T7p29 | gene 5 | DNA polymerase |  |  | 2/154 |  |  |
| 14489 | 137T>C | Leu46Pro | T7p29 | gene 5 | DNA polymerase | 2/161 |  |  |  |  |
| 14494 | 142G>A | Ala48Thr | T7p29 | gene 5 | DNA polymerase |  | 5/156 |  |  |  |
| 14495 | 143C>T | Ala48Val | T7p29 | gene 5 | DNA polymerase | 16/159 |  |  |  |  |
| 14508 | 156A>C | Arg52Arg | T7p29 | gene 5 | DNA polymerase |  | 2/154 | 162/162 | 153/153 | 84/84 |
| 14538 | 186C>A | His62Gln | T7p29 | gene 5 | DNA polymerase |  | 2/174 |  |  |  |
| 14556 | 204A>G | Ala68Ala | T7p29 | gene 5 | DNA polymerase |  |  |  | 2/158 |  |
| 14568 | 216G>T | Leu72Leu | T7p29 | gene 5 | DNA polymerase | 2/151 |  |  |  |  |
| 14629 | 277G>A | Val93Met | T7p29 | gene 5 | DNA polymerase |  |  | 2/199 |  |  |
| 14664 | 312C>T | Asp104Asp | T7p29 | gene 5 | DNA polymerase |  |  |  | 2/162 |  |
| 14674 | 322G>C | Gly108Arg | T7p29 | gene 5 | DNA polymerase |  |  | 2/184 |  |  |
| 14755 | 403A>G | Met135Val | T7p29 | gene 5 | DNA polymerase |  |  |  | 2/151 |  |
| 14776 | 424G>A | Asp142Asn | T7p29 | gene 5 | DNA polymerase | 2/150 |  |  |  |  |
| 14857 | 505G>A | Asp169Asn | T7p29 | gene 5 | DNA polymerase | 6/160 |  |  |  |  |
| 14903 | 551A>G | Lys184Arg | T7p29 | gene 5 | DNA polymerase |  | 2/164 |  |  |  |
| 14935 | 583G>A | Glu195Lys | T7p29 | gene 5 | DNA polymerase |  |  | 3/157 |  |  |
| 15058 | 706A>G | Thr236Ala | T7p29 | gene 5 | DNA polymerase |  |  |  |  | 2/86 |
| 15060 | 708A>G | Thr236Thr | T7p29 | gene 5 | DNA polymerase |  |  |  | 2/155 |  |
| 15125 | 773C>T | Thr258Ile | T7p29 | gene 5 | DNA polymerase |  | 2/154 |  |  |  |
| 15127 | 775G>A | Glu259Lys | T7p29 | gene 5 | DNA polymerase |  | 2/153 |  |  |  |
| **15160** | **808G>A** | **Gly270Ser** | **T7p29** | **gene 5** | **DNA polymerase** | **2/142** |  |  |  |  |
| **15180** | **828T>C** | **His276His** | **T7p29** | **gene 5** | **DNA polymerase** |  |  | **2/146** |  |  |
| **15190** | **838G>A** | **Gly280Ser** | **T7p29** | **gene 5** | **DNA polymerase** |  |  |  | **2/150** |  |
| **15224** | **872C>T** | **Thr291Ile** | **T7p29** | **gene 5** | **DNA polymerase** |  |  | **2/155** |  |  |
| **15314** | **962T>G** | **Val321Gly** | **T7p29** | **gene 5** | **DNA polymerase** | **2/163** |  |  |  |  |
| 15368 | 1016G>C | Arg339Pro | T7p29 | gene 5 | DNA polymerase |  |  |  |  | 2/71 |
| 15371 | 1019A>G | Asp340Gly | T7p29 | gene 5 | DNA polymerase | 3/165 |  |  |  |  |
| 15389 | 1037T>C | Leu346Pro | T7p29 | gene 5 | DNA polymerase |  | 2/150 |  |  |  |
| 15425 | 1073A>C | Asp358Ala | T7p29 | gene 5 | DNA polymerase |  |  |  | 2/139 |  |
| 15449 | 1097A>C | Asp366Ala | T7p29 | gene 5 | DNA polymerase | 155/155 | 149/149 | 167/167 | 129/129 | 81/81 |
| 15517 | 1165T>A | Tyr389Asn | T7p29 | gene 5 | DNA polymerase |  | 2/150 |  |  |  |
| 15583 | 1231G>A | Ala411Thr | T7p29 | gene 5 | DNA polymerase |  |  |  | 2/147 |  |
| 15594 | 1242T>A | Gly414Gly | T7p29 | gene 5 | DNA polymerase | 4/169 |  |  |  |  |
| 15614 | 1262A>G | Asn421Ser | T7p29 | gene 5 | DNA polymerase |  |  |  |  | 2/103 |
| 15631 | 1279A>G | Thr427Ala | T7p29 | gene 5 | DNA polymerase |  |  |  | 2/161 |  |
| 15633 | 1281G>A | Thr427Thr | T7p29 | gene 5 | DNA polymerase |  |  | 2/154 |  |  |
| 15707 | 1355G>A | Arg452His | T7p29 | gene 5 | DNA polymerase |  |  |  | 2/142 |  |
| 15754 | 1402C>G | Pro468Ala | T7p29 | gene 5 | DNA polymerase |  |  | 2/152 |  |  |
| 15762 | 1410T>C | Val470Val | T7p29 | gene 5 | DNA polymerase |  |  |  | 2/136 |  |
| 15838 | 1486T>G | Tyr496Asp | T7p29 | gene 5 | DNA polymerase |  |  | 2/155 |  | 2/102 |
| 15840 | 1488C>A | Tyr496* | T7p29 | gene 5 | DNA polymerase |  | 2/178 | 3/170 | 2/129 |  |
| 15843 | 1491T>C | Ala497Ala | T7p29 | gene 5 | DNA polymerase |  |  | 3/133 |  |  |
| 15845 | 1493A>C | His498Pro | T7p29 | gene 5 | DNA polymerase |  |  | 3/148 |  |  |
| 15848 | 1496A>G | Glu499Gly | T7p29 | gene 5 | DNA polymerase |  | 2/144 |  |  |  |
| 15853 | 1501C>T | Leu501Phe | T7p29 | gene 5 | DNA polymerase |  | 3/183 |  |  |  |
| 15856 | 1504A>T | Asn502Tyr | T7p29 | gene 5 | DNA polymerase | 4/135 |  |  | 4/136 | 3/118 |
| 15858 | 1506C>A | Asn502Lys | T7p29 | gene 5 | DNA polymerase |  |  | 2/167 |  |  |
| 15859 | 1507G>C | Gly503Arg | T7p29 | gene 5 | DNA polymerase | 2/120 |  |  |  | 2/107 |
| 15867 | 1515C>T | Ile505Ile | T7p29 | gene 5 | DNA polymerase |  | 3/181 |  |  |  |
| 15869 | 1517A>T | His506Leu | T7p29 | gene 5 | DNA polymerase |  |  |  | 2/134 |  |
| 15876 | 1524G>A | Lys508Lys | T7p29 | gene 5 | DNA polymerase |  | 4/170 |  | 2/124 | 2/109 |
| 15876 | 1524G>T | Lys508Asn | T7p29 | gene 5 | DNA polymerase |  |  |  | 2/124 |  |
| 15920 | 1568C>A | Thr523Lys | T7p29 | gene 5 | DNA polymerase |  |  |  | 2/130 |  |
| 15922 | 1570T>G | Phe524Val | T7p29 | gene 5 | DNA polymerase |  |  |  | 2/124 |  |
| 15929 | 1577A>G | Tyr526Cys | T7p29 | gene 5 | DNA polymerase | 2/143 |  |  |  |  |
| 15931 | 1579G>T | Gly527Trp | T7p29 | gene 5 | DNA polymerase |  | 4/179 |  |  |  |
| 15946 | 1594G>A | Ala532Thr | T7p29 | gene 5 | DNA polymerase |  |  |  |  | 2/113 |
| 16058 | 1706T>A | Ile569Asn | T7p29 | gene 5 | DNA polymerase |  |  | 2/177 |  |  |
| 16075 | 1723G>A | Glu575Lys | T7p29 | gene 5 | DNA polymerase |  |  | 6/178 |  |  |
| 16091 | 1739T>C | Val580Ala | T7p29 | gene 5 | DNA polymerase |  |  | 2/183 |  |  |
| 16097 | 1745G>A | Gly582Asp | T7p29 | gene 5 | DNA polymerase |  |  |  | 2/136 |  |
| 16099 | 1747G>A | Glu583Lys | T7p29 | gene 5 | DNA polymerase |  |  | 10/179 |  |  |
| 16114 | 1762T>C | Trp588Arg | T7p29 | gene 5 | DNA polymerase |  | 2/185 |  |  |  |
| 16131 | 1779T>A | Ile593Ile | T7p29 | gene 5 | DNA polymerase |  |  |  |  | 2/107 |
| 16270 | 1918C>T | His640Tyr | T7p29 | gene 5 | DNA polymerase | 2/140 |  |  |  |  |
| 16284 | 1932G>A | Gly644Gly | T7p29 | gene 5 | DNA polymerase |  | 5/149 |  |  |  |
| 16307 | 1955T>C | Val652Ala | T7p29 | gene 5 | DNA polymerase |  |  |  | 2/137 |  |
| 16365 | 2013G>C | Glu671Asp | T7p29 | gene 5 | DNA polymerase |  |  |  |  | 2/89 |
| 16371 | 2019A>T | Ala673Ala | T7p29 | gene 5 | DNA polymerase |  | 2/150 |  |  |  |
| 16403 | 2051G>A | Trp684* | T7p29 | gene 5 | DNA polymerase |  |  | 2/123 |  |  |
| 16460 | 2108G>A | Cys703Tyr | T7p29 | gene 5 | DNA polymerase |  |  |  | 2/127 |  |

**EXPERIMENT B**

| **POS** | **NT_CHANGE** | **AA_CHANGE** | **LOCUS_TAG** | **GENE** | **PRODUCT** | **V1** | **V2** | **V3** | **V4** | **V5** |
| --- | --- | --- | --- | --- | --- | --- | --- | --- | --- | --- |
| 3177 | 7A>T | Thr3Ser | T7p07 | gene 1 | T3/T7-like RNA polymerase |  |  |  |  | 2/149 |
| 3232 | 62T>C | Phe21Ser | T7p07 | gene 1 | T3/T7-like RNA polymerase |  |  |  | 2/187 |  |
| 3256 | 86G>T | Gly29Val | T7p07 | gene 1 | T3/T7-like RNA polymerase |  | 2/149 |  |  |  |
| 3268 | 98C>T | Ala33Val | T7p07 | gene 1 | T3/T7-like RNA polymerase |  | 2/150 |  |  |  |
| 3283 | 113C>A | Ala38Asp | T7p07 | gene 1 | T3/T7-like RNA polymerase |  | 2/149 |  |  |  |
| 3284 | 114C>T | Ala38Ala | T7p07 | gene 1 | T3/T7-like RNA polymerase |  | 2/154 |  |  |  |
| 3294 | 124G>T | Glu42* | T7p07 | gene 1 | T3/T7-like RNA polymerase |  | 2/148 |  |  |  |
| 3296 | 126G>A | Glu42Glu | T7p07 | gene 1 | T3/T7-like RNA polymerase |  | 2/154 |  |  |  |
| 3313 | 143A>G | Glu48Gly | T7p07 | gene 1 | T3/T7-like RNA polymerase |  | 2/158 |  |  |  |
| 3363 | 193G>A | Ala65Thr | T7p07 | gene 1 | T3/T7-like RNA polymerase |  | 2/169 |  |  |  |
| 3386 | 216T>G | Pro72Pro | T7p07 | gene 1 | T3/T7-like RNA polymerase |  | 2/170 |  |  |  |
| 3391 | 221T>C | Ile74Thr | T7p07 | gene 1 | T3/T7-like RNA polymerase | 2/142 |  |  |  |  |
| 3407 | 237T>C | Pro79Pro | T7p07 | gene 1 | T3/T7-like RNA polymerase |  |  |  |  | 10/156 |
| 3428 | 258C>G | Asn86Lys | T7p07 | gene 1 | T3/T7-like RNA polymerase |  |  |  | 2/166 |  |
| 3464 | 294G>A | Lys98Lys | T7p07 | gene 1 | T3/T7-like RNA polymerase |  |  |  |  | 2/139 |
| 3495 | 325A>T | Ile109Phe | T7p07 | gene 1 | T3/T7-like RNA polymerase |  |  |  |  | 2/132 |
| 3498 | 328A>G | Lys110Glu | T7p07 | gene 1 | T3/T7-like RNA polymerase |  | 2/160 |  |  |  |
| 3549 | 379A>C | Thr127Pro | T7p07 | gene 1 | T3/T7-like RNA polymerase |  | 159/172 | 120/121 | 174/174 | 142/149 |
| 3587 | 417C>T | Ser139Ser | T7p07 | gene 1 | T3/T7-like RNA polymerase | 2/110 |  |  |  |  |
| 3607 | 437A>G | Glu146Gly | T7p07 | gene 1 | T3/T7-like RNA polymerase | 2/115 |  |  |  |  |
| 3636 | 466G>T | Asp156Tyr | T7p07 | gene 1 | T3/T7-like RNA polymerase |  |  |  | 2/149 |  |
| 3656 | 486C>A | Phe162Leu | T7p07 | gene 1 | T3/T7-like RNA polymerase |  |  |  | 2/165 |  |
| 3693 | 523G>A | Gly175Arg | T7p07 | gene 1 | T3/T7-like RNA polymerase | 9/114 |  |  |  |  |
| 3712 | 542C>T | Ala181Val | T7p07 | gene 1 | T3/T7-like RNA polymerase |  | 3/139 |  |  |  |
| 3713 | 543A>T | Ala181Ala | T7p07 | gene 1 | T3/T7-like RNA polymerase |  | 2/142 |  |  |  |
| 3720 | 550C>A | Gln184Lys | T7p07 | gene 1 | T3/T7-like RNA polymerase |  |  |  | 2/166 |  |
| 3732 | 562G>T | Ala188Ser | T7p07 | gene 1 | T3/T7-like RNA polymerase |  |  |  | 2/164 |  |
| 3733 | 563C>T | Ala188Val | T7p07 | gene 1 | T3/T7-like RNA polymerase |  |  | 2/123 |  |  |
| 3734 | 564T>G | Ala188Ala | T7p07 | gene 1 | T3/T7-like RNA polymerase |  | 2/136 |  |  |  |
| 3739 | 569T>A | Met190Lys | T7p07 | gene 1 | T3/T7-like RNA polymerase |  | 2/135 |  |  |  |
| 3740 | 570G>A | Met190Ile | T7p07 | gene 1 | T3/T7-like RNA polymerase |  |  |  | 2/161 |  |
| 3751 | 581G>A | Gly194Asp | T7p07 | gene 1 | T3/T7-like RNA polymerase |  |  |  |  | 2/167 |
| 3751 | 581G>T | Gly194Asp | T7p07 | gene 1 | T3/T7-like RNA polymerase |  |  |  |  | 2/167 |
| 3753 | 583C>T | Leu195Leu | T7p07 | gene 1 | T3/T7-like RNA polymerase |  |  |  |  | 2/157 |
| 3783 | 613C>T | His205Tyr | T7p07 | gene 1 | T3/T7-like RNA polymerase |  |  | 2/133 |  |  |
| 3783 | 613C>A | His205Asn | T7p07 | gene 1 | T3/T7-like RNA polymerase |  |  |  |  | 2/145 |
| 3796 | 626C>T | Ser209Phe | T7p07 | gene 1 | T3/T7-like RNA polymerase |  |  |  |  | 2/144 |
| 3823 | 653A>G | Glu218Gly | T7p07 | gene 1 | T3/T7-like RNA polymerase |  | 2/138 |  |  |  |
| 3825 | 655A>T | Met219Leu | T7p07 | gene 1 | T3/T7-like RNA polymerase |  |  |  |  | 2/135 |
| 3826 | 656T>G | Met219Arg | T7p07 | gene 1 | T3/T7-like RNA polymerase |  |  | 2/119 |  |  |
| 3828 | 658C>T | Leu220Phe | T7p07 | gene 1 | T3/T7-like RNA polymerase |  | 2/140 |  |  |  |
| 3833 | 663T>G | Ile221Met | T7p07 | gene 1 | T3/T7-like RNA polymerase |  |  |  |  | 2/134 |
| 3834 | 664G>A | Glu222Lys | T7p07 | gene 1 | T3/T7-like RNA polymerase |  |  | 2/127 |  |  |
| 3836 | 666G>T | Glu222Asp | T7p07 | gene 1 | T3/T7-like RNA polymerase |  |  |  | 2/170 |  |
| 3844 | 674G>T | Gly225Val | T7p07 | gene 1 | T3/T7-like RNA polymerase |  | 2/148 |  |  |  |
| 3858 | 688C>T | His230Tyr | T7p07 | gene 1 | T3/T7-like RNA polymerase |  |  |  | 2/162 |  |
| 3886 | 716A>G | Gln239Arg | T7p07 | gene 1 | T3/T7-like RNA polymerase | 2/111 |  |  |  |  |
| 3922 | 752C>A | Ala251Asp | T7p07 | gene 1 | T3/T7-like RNA polymerase |  | 2/161 |  |  |  |
| 3960 | 790A>C | Ile264Leu | T7p07 | gene 1 | T3/T7-like RNA polymerase |  | 2/155 |  |  |  |
| 3967 | 797C>T | Pro266Leu | T7p07 | gene 1 | T3/T7-like RNA polymerase |  |  |  | 2/166 |  |
| 4046 | 876T>C | Arg292Arg | T7p07 | gene 1 | T3/T7-like RNA polymerase |  |  | 2/129 |  |  |
| 4055 | 885G>A | Ala295Ala | T7p07 | gene 1 | T3/T7-like RNA polymerase |  |  | 2/124 |  |  |
| 4069 | 899A>G | His300Arg | T7p07 | gene 1 | T3/T7-like RNA polymerase |  | 2/142 |  |  |  |
| 4093 | 923A>T | Tyr308Phe | T7p07 | gene 1 | T3/T7-like RNA polymerase |  | 2/154 |  |  |  |
| 4101 | 931G>A | Val311Ile | T7p07 | gene 1 | T3/T7-like RNA polymerase |  |  |  | 4/174 |  |
| 4111 | 941C>T | Pro314Leu | T7p07 | gene 1 | T3/T7-like RNA polymerase |  | 2/167 |  |  |  |
| 4125 | 955G>T | Ala319Ser | T7p07 | gene 1 | T3/T7-like RNA polymerase |  | 3/165 |  |  |  |
| 4133 | 963C>T | Asn321Asn | T7p07 | gene 1 | T3/T7-like RNA polymerase |  | 2/170 |  |  |  |
| 4177 | 1007C>T | Ala336Val | T7p07 | gene 1 | T3/T7-like RNA polymerase |  |  | 2/164 |  |  |
| 4205 | 1035G>A | Lys345Lys | T7p07 | gene 1 | T3/T7-like RNA polymerase |  | 2/189 |  |  |  |
| 4222 | 1052A>G | Asp351Gly | T7p07 | gene 1 | T3/T7-like RNA polymerase |  |  |  |  | 2/156 |
| 4224 | 1054A>G | Ile352Val | T7p07 | gene 1 | T3/T7-like RNA polymerase |  |  | 2/166 |  |  |
| 4235 | 1065T>C | Ile355Ile | T7p07 | gene 1 | T3/T7-like RNA polymerase |  |  | 2/172 |  |  |
| 4258 | 1088A>G | Lys363Arg | T7p07 | gene 1 | T3/T7-like RNA polymerase |  |  |  |  | 2/154 |
| 4263 | 1093G>C | Glu365Gln | T7p07 | gene 1 | T3/T7-like RNA polymerase |  | 2/181 |  |  |  |
| 4265 | 1095A>G | Glu365Glu | T7p07 | gene 1 | T3/T7-like RNA polymerase |  | 3/176 |  |  |  |
| 4289 | 1119T>G | Ala373Ala | T7p07 | gene 1 | T3/T7-like RNA polymerase |  | 2/170 |  |  |  |
| 4289 | 1119T>C | Ala373Ala | T7p07 | gene 1 | T3/T7-like RNA polymerase |  |  |  |  | 2/142 |
| 4291 | 1121T>C | Leu374Pro | T7p07 | gene 1 | T3/T7-like RNA polymerase |  | 2/167 |  |  |  |
| 4315 | 1145C>T | Ala382Val | T7p07 | gene 1 | T3/T7-like RNA polymerase |  |  |  |  | 2/146 |
| 4325 | 1155C>A | Tyr385* | T7p07 | gene 1 | T3/T7-like RNA polymerase |  | 2/179 |  |  |  |
| 4337 | 1167G>A | Lys389Lys | T7p07 | gene 1 | T3/T7-like RNA polymerase |  | 2/157 |  |  |  |
| 4366 | 1196A>C | Glu399Ala | T7p07 | gene 1 | T3/T7-like RNA polymerase |  | 2/161 |  |  |  |
| 4372 | 1202T>A | Met401Lys | T7p07 | gene 1 | T3/T7-like RNA polymerase |  | 2/158 |  |  |  |
| 4392 | 1222T>C | Phe408Leu | T7p07 | gene 1 | T3/T7-like RNA polymerase |  | 2/147 |  |  |  |
| 4400 | 1230C>T | Asn410Asn | T7p07 | gene 1 | T3/T7-like RNA polymerase |  |  |  | 2/188 |  |
| 4411 | 1241T>C | Ile414Thr | T7p07 | gene 1 | T3/T7-like RNA polymerase |  |  |  | 2/176 |  |
| 4456 | 1286T>A | Val429Glu | T7p07 | gene 1 | T3/T7-like RNA polymerase |  | 2/133 |  |  |  |
| 4459 | 1289C>T | Ser430Leu | T7p07 | gene 1 | T3/T7-like RNA polymerase |  | 2/140 |  |  |  |
| 4464 | 1294T>C | Phe432Leu | T7p07 | gene 1 | T3/T7-like RNA polymerase |  |  |  |  | 2/132 |
| 4475 | 1305A>T | Gln435His | T7p07 | gene 1 | T3/T7-like RNA polymerase |  |  |  |  | 2/132 |
| 4612 | 1442T>G | Phe481Cys | T7p07 | gene 1 | T3/T7-like RNA polymerase |  | 2/154 |  |  |  |
| 4658 | 1488A>T | Pro496Pro | T7p07 | gene 1 | T3/T7-like RNA polymerase |  | 3/151 |  |  |  |
| 4663 | 1493A>G | Glu498Gly | T7p07 | gene 1 | T3/T7-like RNA polymerase | 2/119 |  |  |  |  |
| 4681 | 1511A>G | Glu504Gly | T7p07 | gene 1 | T3/T7-like RNA polymerase |  | 2/139 |  |  |  |
| 4717 | 1547T>C | Phe516Ser | T7p07 | gene 1 | T3/T7-like RNA polymerase |  |  |  |  | 2/129 |
| 4719 | 1549G>A | Glu517Lys | T7p07 | gene 1 | T3/T7-like RNA polymerase |  |  |  |  | 2/131 |
| 4732 | 1562T>G | Val521Gly | T7p07 | gene 1 | T3/T7-like RNA polymerase |  |  |  |  | 2/132 |
| 4749 | 1579A>G | Ser527Gly | T7p07 | gene 1 | T3/T7-like RNA polymerase |  | 2/138 |  |  |  |
| 4882 | 1712A>G | Tyr571Cys | T7p07 | gene 1 | T3/T7-like RNA polymerase | 2/174 |  | 2/140 |  |  |
| 4904 | 1734C>A | Val578Val | T7p07 | gene 1 | T3/T7-like RNA polymerase | 2/169 |  |  |  |  |
| 4950 | 1780G>A | Val594Ile | T7p07 | gene 1 | T3/T7-like RNA polymerase |  | 3/152 |  |  |  |
| 4959 | 1789G>A | Val597Met | T7p07 | gene 1 | T3/T7-like RNA polymerase |  | 2/157 |  |  |  |
| 4977 | 1807G>A | Gly603Ser | T7p07 | gene 1 | T3/T7-like RNA polymerase |  |  | 5/135 | 181/185 | 145/148 |
| 4979 | 1809T>A | Gly603Gly | T7p07 | gene 1 | T3/T7-like RNA polymerase |  |  |  | 2/181 |  |
| 4990 | 1820A>G | Glu607Gly | T7p07 | gene 1 | T3/T7-like RNA polymerase |  |  |  |  | 2/138 |
| 5016 | 1846C>T | Leu616Leu | T7p07 | gene 1 | T3/T7-like RNA polymerase |  | 2/144 |  |  |  |
| 5017 | 1847T>G | Leu616Arg | T7p07 | gene 1 | T3/T7-like RNA polymerase |  | 2/141 |  |  |  |
| 5044 | 1874T>G | Val625Gly | T7p07 | gene 1 | T3/T7-like RNA polymerase |  | 2/143 |  |  |  |
| 5064 | 1894C>T | Arg632Cys | T7p07 | gene 1 | T3/T7-like RNA polymerase |  |  |  | 2/176 |  |
| 5071 | 1901T>G | Val634Gly | T7p07 | gene 1 | T3/T7-like RNA polymerase |  | 2/153 |  |  |  |
| 5134 | 1964T>A | Ile655Asn | T7p07 | gene 1 | T3/T7-like RNA polymerase |  | 2/158 |  |  |  |
| 5136 | 1966C>T | Gln656* | T7p07 | gene 1 | T3/T7-like RNA polymerase |  | 3/157 |  |  |  |
| 5166 | 1996A>G | Met666Val | T7p07 | gene 1 | T3/T7-like RNA polymerase |  |  | 2/124 |  |  |
| 5170 | 2000T>C | Phe667Ser | T7p07 | gene 1 | T3/T7-like RNA polymerase |  | 2/163 |  |  |  |
| 5224 | 2054T>A | Val685Glu | T7p07 | gene 1 | T3/T7-like RNA polymerase |  | 2/164 |  |  |  |
| 5233 | 2063C>T | Thr688Met | T7p07 | gene 1 | T3/T7-like RNA polymerase |  |  | 6/125 |  |  |
| 5254 | 2084C>T | Ala695Val | T7p07 | gene 1 | T3/T7-like RNA polymerase |  | 4/169 |  |  |  |
| 5263 | 2093G>A | Trp698* | T7p07 | gene 1 | T3/T7-like RNA polymerase | 2/145 |  |  |  |  |
| 5277 | 2107G>A | Ala703Thr | T7p07 | gene 1 | T3/T7-like RNA polymerase |  | 2/171 |  |  |  |
| 5278 | 2108C>A | Ala703Asp | T7p07 | gene 1 | T3/T7-like RNA polymerase |  | 2/171 |  |  |  |
| 5317 | 2147G>T | Gly716Val | T7p07 | gene 1 | T3/T7-like RNA polymerase | 2/129 |  |  |  |  |
| 5336 | 2166T>C | Arg722Arg | T7p07 | gene 1 | T3/T7-like RNA polymerase | 3/128 |  |  |  |  |
| 5340 | 2170G>A | Ala724Thr | T7p07 | gene 1 | T3/T7-like RNA polymerase |  | 2/157 |  |  |  |
| 5363 | 2193T>C | Asp731Asp | T7p07 | gene 1 | T3/T7-like RNA polymerase | 2/132 |  |  |  |  |
| 5367 | 2197T>G | Phe733Val | T7p07 | gene 1 | T3/T7-like RNA polymerase |  | 2/150 |  |  |  |
| 5368 | 2198T>A | Phe733Tyr | T7p07 | gene 1 | T3/T7-like RNA polymerase |  | 2/149 |  |  |  |
| 5369 | 2199C>A | Phe733Leu | T7p07 | gene 1 | T3/T7-like RNA polymerase |  | 2/152 |  |  |  |
| 5386 | 2216A>T | Tyr739Phe | T7p07 | gene 1 | T3/T7-like RNA polymerase |  | 2/150 |  |  |  |
| 5406 | 2236C>T | Arg746Cys | T7p07 | gene 1 | T3/T7-like RNA polymerase | 2/131 |  | 2/116 |  |  |
| 5408 | 2238C>T | Arg746Arg | T7p07 | gene 1 | T3/T7-like RNA polymerase |  |  | 2/113 |  |  |
| 5421 | 2251T>G | Phe751Val | T7p07 | gene 1 | T3/T7-like RNA polymerase | 2/125 |  |  |  |  |
| 5446 | 2276C>T | Pro759Leu | T7p07 | gene 1 | T3/T7-like RNA polymerase |  |  | 2/115 |  |  |
| 5474 | 2304G>A | Glu768Glu | T7p07 | gene 1 | T3/T7-like RNA polymerase |  | 2/147 |  |  |  |
| 5485 | 2315A>G | His772Arg | T7p07 | gene 1 | T3/T7-like RNA polymerase |  |  |  |  | 2/143 |
| 5548 | 2378A>C | Lys793Thr | T7p07 | gene 1 | T3/T7-like RNA polymerase |  |  |  |  | 2/147 |
| 5597 | 2427G>A | Leu809Leu | T7p07 | gene 1 | T3/T7-like RNA polymerase |  |  |  | 2/176 |  |
| 5598 | 2428A>G | Ile810Val | T7p07 | gene 1 | T3/T7-like RNA polymerase |  | 3/135 |  |  |  |
| 5603 | 2433C>T | His811His | T7p07 | gene 1 | T3/T7-like RNA polymerase |  |  |  | 2/192 |  |
| 5660 | 2490A>G | Glu830Glu | T7p07 | gene 1 | T3/T7-like RNA polymerase |  |  | 2/106 |  |  |
| 5713 | 2543A>G | Gln848Arg | T7p07 | gene 1 | T3/T7-like RNA polymerase |  |  |  |  | 2/150 |
| 5719 | 2549C>A | Ala850Asp | T7p07 | gene 1 | T3/T7-like RNA polymerase |  |  | 24/106 | 3/212 |  |
| 5728 | 2558T>C | Leu853Ser | T7p07 | gene 1 | T3/T7-like RNA polymerase | 2/142 |  |  |  |  |
| 5745 | 2575G>T | Asp859Tyr | T7p07 | gene 1 | T3/T7-like RNA polymerase |  |  |  |  | 2/155 |
| 5759 | 2589A>G | Ala863Ala | T7p07 | gene 1 | T3/T7-like RNA polymerase |  | 2/148 |  |  |  |
| 5804 | 2634G>A | Ser878Ser | T7p07 | gene 1 | T3/T7-like RNA polymerase |  |  |  | 2/192 |  |
| 5810 | 2640C>T | Phe880Phe | T7p07 | gene 1 | T3/T7-like RNA polymerase | 2/151 |  |  |  |  |
| 14359 | 7G>A | Val3Ile | T7p29 | gene 5 | DNA polymerase |  |  | 5/143 | 5/186 |  |
| 14359 | 7G>T |  | T7p29 | gene 5 | DNA polymerase |  |  |  | 2/186 |  |
| 14389 | 37G>C | Glu13Gln | T7p29 | gene 5 | DNA polymerase | 2/156 |  |  |  |  |
| 14392 | 40A>G | Ser14Gly | T7p29 | gene 5 | DNA polymerase |  |  |  | 2/193 |  |
| 14399 | 47C>A | Thr16Asn | T7p29 | gene 5 | DNA polymerase |  |  | 2/149 | 10/199 |  |
| 14406 | 54C>T | Phe18Phe | T7p29 | gene 5 | DNA polymerase |  | 2/174 |  |  |  |
| 14419 | 67A>G | Ile23Val | T7p29 | gene 5 | DNA polymerase |  | 12/182 |  |  |  |
| 14424 | 72C>T | Tyr24Tyr | T7p29 | gene 5 | DNA polymerase | 145/145 | 183/183 | 160/160 | 211/213 | 149/149 |
| 14440 | 88G>A | Glu30Lys | T7p29 | gene 5 | DNA polymerase |  |  | 3/150 |  |  |
| 14461 | 109A>G | Ser37Gly | T7p29 | gene 5 | DNA polymerase |  |  |  |  | 2/141 |
| 14462 | 110G>A | Ser37Asn | T7p29 | gene 5 | DNA polymerase |  |  | 2/138 | 2/192 |  |
| 14467 | 115T>C | Phe39Leu | T7p29 | gene 5 | DNA polymerase |  |  |  | 2/189 |  |
| 14495 | 143C>T | Ala48Val | T7p29 | gene 5 | DNA polymerase |  |  |  | 2/176 |  |
| 14508 | 156A>C | Arg52Arg | T7p29 | gene 5 | DNA polymerase |  |  |  | 2/170 |  |
| 14511 | 159C>A | Gly53Gly | T7p29 | gene 5 | DNA polymerase |  |  | 3/125 |  |  |
| 14522 | 170T>C | Val57Ala | T7p29 | gene 5 | DNA polymerase |  |  | 2/127 |  |  |
| 14620 | 268G>T | Asp90Tyr | T7p29 | gene 5 | DNA polymerase |  |  |  |  | 2/135 |
| 14623 | 271A>T | Thr91Ser | T7p29 | gene 5 | DNA polymerase |  |  |  |  | 2/136 |
| 14662 | 310G>A | Asp104Asn | T7p29 | gene 5 | DNA polymerase | 2/167 |  |  |  |  |
| 14680 | 328C>A | Leu110Met | T7p29 | gene 5 | DNA polymerase |  | 2/148 |  |  |  |
| 14693 | 341A>G | Lys114Arg | T7p29 | gene 5 | DNA polymerase |  | 2/145 |  |  |  |
| 14751 | 399C>T | Gly133Gly | T7p29 | gene 5 | DNA polymerase |  |  |  |  | 2/109 |
| 14753 | 401A>G | Glu134Gly | T7p29 | gene 5 | DNA polymerase | 2/170 |  |  |  |  |
| 14773 | 421G>A | Asp141Asn | T7p29 | gene 5 | DNA polymerase |  |  |  | 2/146 |  |
| 14786 | 434G>A | Arg145His | T7p29 | gene 5 | DNA polymerase |  |  |  |  | 2/108 |
| 14819 | 467A>G | Asp156Gly | T7p29 | gene 5 | DNA polymerase |  |  | 2/136 |  |  |
| 14898 | 546T>C | Leu182Leu | T7p29 | gene 5 | DNA polymerase |  |  |  | 2/143 |  |
| 14961 | 609C>G | Tyr203* | T7p29 | gene 5 | DNA polymerase |  |  |  | 2/152 |  |
| 14996 | 644A>T | Asp215Val | T7p29 | gene 5 | DNA polymerase |  | 2/137 |  |  |  |
| 15002 | 650A>G | Glu217Gly | T7p29 | gene 5 | DNA polymerase |  |  |  | 2/150 |  |
| 15029 | 677A>C | Lys226Thr | T7p29 | gene 5 | DNA polymerase |  | 2/143 |  |  |  |
| 15081 | 729C>G | Tyr243* | T7p29 | gene 5 | DNA polymerase |  | 2/151 |  |  |  |
| 15092 | 740C>A | Ala247Asp | T7p29 | gene 5 | DNA polymerase |  | 2/147 |  |  |  |
| 15093 | 741T>C | Ala247Ala | T7p29 | gene 5 | DNA polymerase |  |  |  | 2/166 |  |
| 15096 | 744T>G | Ala248Ala | T7p29 | gene 5 | DNA polymerase |  |  | 6/125 |  |  |
| 15099 | 747C>G | Arg249Arg | T7p29 | gene 5 | DNA polymerase |  | 2/142 |  |  |  |
| **15152** | **800C>T** | **Pro267Leu** | **T7p29** | **gene 5** | **DNA polymerase** |  |  |  | **2/152** |  |
| **15209** | **857A>T** | **Tyr286Phe** | **T7p29** | **gene 5** | **DNA polymerase** |  |  |  | **2/155** |  |
| **15240** | **888T>A** | **Gly296Gly** | **T7p29** | **gene 5** | **DNA polymerase** |  |  |  | **2/142** |  |
| **15242** | **890T>G** | **Ile297Ser** | **T7p29** | **gene 5** | **DNA polymerase** | **2/119** |  |  |  |  |
| **15326** | **974C>T** | **Pro325Leu** | **T7p29** | **gene 5** | **DNA polymerase** |  |  | **2/131** |  |  |
| **15329** | **977A>G** | **Tyr326Cys** | **T7p29** | **gene 5** | **DNA polymerase** |  |  |  | **2/148** |  |
| **15361** | **1009T>C** | **Ser337Pro** | **T7p29** | **gene 5** | **DNA polymerase** |  |  |  |  | **2/125** |
| 15380 | 1028A>G | Gln343Arg | T7p29 | gene 5 | DNA polymerase |  |  | 26/124 | 11/146 |  |
| 15383 | 1031A>G | Lys344Arg | T7p29 | gene 5 | DNA polymerase | 2/137 |  |  |  |  |
| 15391 | 1039C>A | Gln347Lys | T7p29 | gene 5 | DNA polymerase |  | 2/140 |  |  |  |
| 15405 | 1053G>A | Trp351* | T7p29 | gene 5 | DNA polymerase | 2/139 |  |  |  |  |
| 15418 | 1066T>C | Tyr356His | T7p29 | gene 5 | DNA polymerase |  | 2/138 |  |  | 2/132 |
| 15419 | 1067A>G | Tyr356Cys | T7p29 | gene 5 | DNA polymerase |  |  |  | 2/139 |  |
| 15445 | 1093G>A | Asp365Asn | T7p29 | gene 5 | DNA polymerase |  |  |  | 9/137 |  |
| 15494 | 1142C>T | Ala381Val | T7p29 | gene 5 | DNA polymerase |  |  |  | 2/142 |  |
| 15501 | 1149C>G | Ile383Met | T7p29 | gene 5 | DNA polymerase | 2/154 |  |  |  |  |
| 15512 | 1160A>G | Lys387Arg | T7p29 | gene 5 | DNA polymerase |  |  | 6/140 |  |  |
| 15529 | 1177C>A | Gln393Lys | T7p29 | gene 5 | DNA polymerase | 2/151 |  |  |  |  |
| 15533 | 1181A>G | Lys394Arg | T7p29 | gene 5 | DNA polymerase | 147/147 | 133/133 | 139/139 | 154/154 | 126/126 |
| 15575 | 1223G>A | Arg408His | T7p29 | gene 5 | DNA polymerase |  |  |  | 2/160 |  |
| 15581 | 1229T>A | Val410Asp | T7p29 | gene 5 | DNA polymerase |  |  |  |  | 2/122 |
| 15592 | 1240G>A | Gly414Ser | T7p29 | gene 5 | DNA polymerase |  |  |  |  | 2/124 |
| 15692 | 1340A>G | Tyr447Cys | T7p29 | gene 5 | DNA polymerase |  |  | 2/168 |  |  |
| 15702 | 1350G>A | Gln450Gln | T7p29 | gene 5 | DNA polymerase |  |  | 2/161 |  |  |
| 15708 | 1356C>T | Arg452Arg | T7p29 | gene 5 | DNA polymerase |  |  |  | 5/164 |  |
| 15784 | 1432G>A | Gly478Ser | T7p29 | gene 5 | DNA polymerase |  |  |  |  | 2/127 |
| 15814 | 1462A>T | Met488Leu | T7p29 | gene 5 | DNA polymerase | 2/142 |  |  |  |  |
| 15840 | 1488C>T | Tyr496Tyr | T7p29 | gene 5 | DNA polymerase |  |  |  |  | 2/131 |
| 15843 | 1491T>C | Ala497Ala | T7p29 | gene 5 | DNA polymerase |  |  | 4/129 |  |  |
| 15845 | 1493A>C | His498Pro | T7p29 | gene 5 | DNA polymerase |  |  | 2/143 |  |  |
| 15846 | 1494C>A | His498Gln | T7p29 | gene 5 | DNA polymerase |  | 2/142 |  |  |  |
| 15848 | 1496A>G | Glu499Gly | T7p29 | gene 5 | DNA polymerase |  | 2/119 |  | 2/121 |  |
| 15850 | 1498A>G | Ile500Val | T7p29 | gene 5 | DNA polymerase | 3/123 |  |  |  |  |
| 15853 | 1501C>T | Leu501Phe | T7p29 | gene 5 | DNA polymerase | 2/147 | 2/142 |  |  |  |
| 15853 | 1501C>A | Leu501Ile | T7p29 | gene 5 | DNA polymerase |  |  |  | 2/166 |  |
| 15854 | 1502T>C | Leu501Pro | T7p29 | gene 5 | DNA polymerase |  | 2/140 |  | 3/168 |  |
| 15856 | 1504A>T | Asn502Tyr | T7p29 | gene 5 | DNA polymerase |  |  |  |  | 3/132 |
| 15858 | 1506C>A | Asn502Lys | T7p29 | gene 5 | DNA polymerase |  |  |  | 2/162 |  |
| 15867 | 1515C>T | Ile505Ile | T7p29 | gene 5 | DNA polymerase |  |  |  | 2/160 |  |
| 15869 | 1517A>C | His506Pro | T7p29 | gene 5 | DNA polymerase |  |  |  | 2/160 |  |
| 15876 | 1524G>A | Lys508Lys | T7p29 | gene 5 | DNA polymerase |  |  | 3/152 | 4/157 |  |
| 15879 | 1527C>A | Asn509Lys | T7p29 | gene 5 | DNA polymerase |  | 2/121 | 2/150 |  | 2/123 |
| 15895 | 1543C>A | Leu515Ile | T7p29 | gene 5 | DNA polymerase |  |  |  | 3/150 |  |
| 15897 | 1545A>T | Leu515Leu | T7p29 | gene 5 | DNA polymerase | 2/141 | 2/127 |  |  |  |
| 15902 | 1550C>A | Thr517Asn | T7p29 | gene 5 | DNA polymerase | 2/136 |  |  |  |  |
| 15918 | 1566G>A | Lys522Lys | T7p29 | gene 5 | DNA polymerase | 2/142 | 2/132 | 2/140 | 2/155 |  |
| 15926 | 1574T>A | Ile525Asn | T7p29 | gene 5 | DNA polymerase |  |  |  | 2/170 |  |
| 15932 | 1580G>T | Gly527Val | T7p29 | gene 5 | DNA polymerase | 2/147 |  |  |  |  |
| 15945 | 1593T>A | Gly531Gly | T7p29 | gene 5 | DNA polymerase |  |  |  | 2/173 |  |
| 15968 | 1616A>T | Gln539Leu | T7p29 | gene 5 | DNA polymerase |  | 2/132 |  |  |  |
| 15973 | 1621G>T | Val541Phe | T7p29 | gene 5 | DNA polymerase |  |  |  |  | 2/145 |
| 16012 | 1660A>G | Lys554Glu | T7p29 | gene 5 | DNA polymerase |  |  | 2/107 |  |  |
| 16045 | 1693C>A | Leu565Ile | T7p29 | gene 5 | DNA polymerase |  | 2/135 |  |  |  |
| 16067 | 1715C>T | Thr572Ile | T7p29 | gene 5 | DNA polymerase |  | 2/137 |  |  |  |
| 16075 | 1723G>A | Glu575Lys | T7p29 | gene 5 | DNA polymerase |  |  | 6/102 |  |  |
| 16080 | 1728C>T | Ser576Ser | T7p29 | gene 5 | DNA polymerase | 2/129 |  |  |  |  |
| 16099 | 1747G>A | Glu583Lys | T7p29 | gene 5 | DNA polymerase |  |  |  | 5/202 |  |
| 16109 | 1757T>A | Val586Asp | T7p29 | gene 5 | DNA polymerase |  | 2/143 |  |  |  |
| 16156 | 1804C>T | His602Tyr | T7p29 | gene 5 | DNA polymerase |  |  | 4/98 |  |  |
| 16239 | 1887C>T | Thr629Thr | T7p29 | gene 5 | DNA polymerase |  |  |  | 2/180 |  |
| 16245 | 1893G>T | Glu631Asp | T7p29 | gene 5 | DNA polymerase |  |  |  |  | 2/94 |
| 16251 | 1899C>T | Leu633Leu | T7p29 | gene 5 | DNA polymerase | 3/110 |  |  |  |  |
| 16290 | 1938T>A | Phe646Leu | T7p29 | gene 5 | DNA polymerase |  | 2/159 |  |  |  |
| 16293 | 1941G>A | Ala647Ala | T7p29 | gene 5 | DNA polymerase |  | 2/155 |  |  |  |
| 16393 | 2041G>A | Gly681Arg | T7p29 | gene 5 | DNA polymerase |  |  |  |  | 2/92 |
| 16425 | 2073T>C | Asp691Asp | T7p29 | gene 5 | DNA polymerase |  |  |  |  | 2/87 |
| 16428 | 2076C>T | Thr692Thr | T7p29 | gene 5 | DNA polymerase |  | 2/169 |  |  |  |
| 16453 | 2101G>T | Ala701Ser | T7p29 | gene 5 | DNA polymerase |  | 2/178 |  |  |  |
| 16456 | 2104A>G | Ile702Val | T7p29 | gene 5 | DNA polymerase |  |  | 2/140 |  |  |
| 16463 | 2111A>C | His704Pro | T7p29 | gene 5 | DNA polymerase | 2/133 | 2/182 |  |  |  |
| 16463 | 2111A>G |  | T7p29 | gene 5 | DNA polymerase |  | 2/182 |  |  |  |

**EXPERIMENT C**

| **POS** | **NT_CHANGE** | **AA_CHANGE** | **LOCUS_TAG** | **GENE** | **PRODUCT** | **V1** | **V2** | **V3** | **V4** | **V5** |
| --- | --- | --- | --- | --- | --- | --- | --- | --- | --- | --- |
| 3206 | 36T>G | Ser12Ser | T7p07 | gene 1 | T3/T7-like RNA polymerase |  | 2/195 |  |  |  |
| 3262 | 92G>A | Arg31His | T7p07 | gene 1 | T3/T7-like RNA polymerase |  |  |  | 237/241 | 222/222 |
| 3306 | 136A>C | Met46Leu | T7p07 | gene 1 | T3/T7-like RNA polymerase |  |  | 4/221 |  |  |
| 3418 | 248C>T | Ala83Val | T7p07 | gene 1 | T3/T7-like RNA polymerase |  |  | 2/196 |  |  |
| 3494 | 324A>G | Glu108Glu | T7p07 | gene 1 | T3/T7-like RNA polymerase |  | 6/189 |  |  |  |
| 3504 | 334G>A | Glu112Lys | T7p07 | gene 1 | T3/T7-like RNA polymerase |  | 2/189 |  |  |  |
| 3518 | 348C>G | Tyr116* | T7p07 | gene 1 | T3/T7-like RNA polymerase | 2/176 |  |  |  |  |
| 3626 | 456T>C | Gly152Gly | T7p07 | gene 1 | T3/T7-like RNA polymerase |  | 7/220 |  |  |  |
| 3651 | 481C>T | His161Tyr | T7p07 | gene 1 | T3/T7-like RNA polymerase | 105/199 | 222/222 | 219/222 | 275/278 | 232/241 |
| 3668 | 498T>G | Val166Val | T7p07 | gene 1 | T3/T7-like RNA polymerase | 2/190 |  |  |  |  |
| 3684 | 514A>G | Lys172Glu | T7p07 | gene 1 | T3/T7-like RNA polymerase | 2/186 |  |  |  |  |
| 3708 | 538A>G | Lys180Glu | T7p07 | gene 1 | T3/T7-like RNA polymerase | 99/185 | 217/217 | 217/224 | 267/270 | 229/240 |
| 3849 | 679G>T | Val227Phe | T7p07 | gene 1 | T3/T7-like RNA polymerase |  |  |  |  | 3/244 |
| 3866 | 696A>G | Gln232Gln | T7p07 | gene 1 | T3/T7-like RNA polymerase | 2/182 |  |  |  |  |
| 3896 | 726G>A | Glu242Glu | T7p07 | gene 1 | T3/T7-like RNA polymerase | 2/180 |  |  |  |  |
| 4046 | 876T>C | Arg292Arg | T7p07 | gene 1 | T3/T7-like RNA polymerase | 2/141 |  |  |  |  |
| 4055 | 885G>C | Ala295Ala | T7p07 | gene 1 | T3/T7-like RNA polymerase | 2/137 |  |  |  |  |
| 4109 | 939G>A | Met313Ile | T7p07 | gene 1 | T3/T7-like RNA polymerase |  | 17/179 | 223/224 | 239/247 | 281/285 |
| 4330 | 1160A>G | Lys387Arg | T7p07 | gene 1 | T3/T7-like RNA polymerase | 2/177 |  |  |  |  |
| 4343 | 1173C>A | Arg391Arg | T7p07 | gene 1 | T3/T7-like RNA polymerase |  |  | 2/183 |  |  |
| 4370 | 1200C>T | Phe400Phe | T7p07 | gene 1 | T3/T7-like RNA polymerase |  |  | 2/189 |  |  |
| 4404 | 1234A>G | Lys412Glu | T7p07 | gene 1 | T3/T7-like RNA polymerase |  |  |  |  | 10/257 |
| 4419 | 1249C>T | Pro417Ser | T7p07 | gene 1 | T3/T7-like RNA polymerase |  |  |  |  | 3/250 |
| 4442 | 1272T>C | Gly424Gly | T7p07 | gene 1 | T3/T7-like RNA polymerase | 2/179 |  |  |  |  |
| 4468 | 1298A>T | Asn433Ile | T7p07 | gene 1 | T3/T7-like RNA polymerase | 2/174 |  |  |  |  |
| 4490 | 1320C>T | Thr440Thr | T7p07 | gene 1 | T3/T7-like RNA polymerase | 2/175 |  |  |  |  |
| 4582 | 1412A>G | Asp471Gly | T7p07 | gene 1 | T3/T7-like RNA polymerase |  | 2/199 |  |  |  |
| 4701 | 1531T>C | Phe511Leu | T7p07 | gene 1 | T3/T7-like RNA polymerase |  | 2/178 |  |  |  |
| 4708 | 1538C>T | Ala513Val | T7p07 | gene 1 | T3/T7-like RNA polymerase |  | 2/179 |  |  |  |
| 4802 | 1632G>A | Gln544Gln | T7p07 | gene 1 | T3/T7-like RNA polymerase |  | 2/176 |  |  |  |
| 4831 | 1661T>A | Val554Glu | T7p07 | gene 1 | T3/T7-like RNA polymerase |  | 2/194 |  |  |  |
| 4888 | 1718T>C | Ile573Thr | T7p07 | gene 1 | T3/T7-like RNA polymerase |  |  |  | 48/262 |  |
| 5036 | 1866T>C | Ala622Ala | T7p07 | gene 1 | T3/T7-like RNA polymerase |  |  |  |  | 4/274 |
| 5057 | 1887G>A | Val629Val | T7p07 | gene 1 | T3/T7-like RNA polymerase |  |  |  | 10/211 |  |
| 5273 | 2103T>G | Ser701Ser | T7p07 | gene 1 | T3/T7-like RNA polymerase |  | 149/208 | 239/240 | 226/226 | 269/279 |
| 5355 | 2185A>G | Thr729Ala | T7p07 | gene 1 | T3/T7-like RNA polymerase |  |  |  | 3/215 |  |
| 5418 | 2248A>G | Met750Val | T7p07 | gene 1 | T3/T7-like RNA polymerase |  | 211/211 | 221/222 | 221/224 | 224/228 |
| 5539 | 2369A>G | His790Arg | T7p07 | gene 1 | T3/T7-like RNA polymerase |  | 2/189 |  |  |  |
| 5581 | 2411T>C | Ile804Thr | T7p07 | gene 1 | T3/T7-like RNA polymerase | 2/175 |  |  |  |  |
| 5620 | 2450T>C | Ile817Thr | T7p07 | gene 1 | T3/T7-like RNA polymerase |  |  | 2/188 |  |  |
| 5668 | 2498T>C | Val833Ala | T7p07 | gene 1 | T3/T7-like RNA polymerase |  |  | 4/188 |  |  |
| 5710 | 2540A>G | Asp847Gly | T7p07 | gene 1 | T3/T7-like RNA polymerase | 2/189 |  |  |  |  |
| 5759 | 2589A>G | Ala863Ala | T7p07 | gene 1 | T3/T7-like RNA polymerase | 2/181 |  |  | 3/236 |  |
| 5774 | 2604T>A | Gly868Gly | T7p07 | gene 1 | T3/T7-like RNA polymerase |  | 2/185 |  |  |  |
| 5788 | 2618G>A | Arg873His | T7p07 | gene 1 | T3/T7-like RNA polymerase |  |  |  |  | 3/271 |
| 5800 | 2630A>G | Glu877Gly | T7p07 | gene 1 | T3/T7-like RNA polymerase | 2/170 |  |  |  |  |
| 14374 | 22G>A | Ala8Thr | T7p29 | gene 5 | DNA polymerase |  | 25/191 |  |  |  |
| 14390 | 38A>G | Glu13Gly | T7p29 | gene 5 | DNA polymerase |  |  | 4/234 |  |  |
| 14395 | 43G>A | Val15Ile | T7p29 | gene 5 | DNA polymerase |  | 2/199 |  |  |  |
| 14403 | 51G>A | Lys17Lys | T7p29 | gene 5 | DNA polymerase |  | 7/202 |  |  |  |
| 14440 | 88G>A | Glu30Lys | T7p29 | gene 5 | DNA polymerase |  | 88/199 |  |  |  |
| 14442 | 90G>A | Glu30Glu | T7p29 | gene 5 | DNA polymerase |  | 13/197 |  |  |  |
| 14446 | 94G>A | Val32Ile | T7p29 | gene 5 | DNA polymerase |  | 36/203 | 219/219 | 239/239 | 267/268 |
| 14464 | 112G>A | Asp38Asn | T7p29 | gene 5 | DNA polymerase |  | 15/195 |  |  |  |
| 14479 | 127C>T | Leu43Leu | T7p29 | gene 5 | DNA polymerase |  | 4/187 |  |  |  |
| 14505 | 153A>G | Ala51Ala | T7p29 | gene 5 | DNA polymerase |  | 2/179 |  |  |  |
| 14520 | 168T>A | Ile56Ile | T7p29 | gene 5 | DNA polymerase |  | 2/194 |  |  |  |
| 14523 | 171G>A | Val57Val | T7p29 | gene 5 | DNA polymerase |  |  |  | 3/209 |  |
| 14561 | 209C>T | Thr70Ile | T7p29 | gene 5 | DNA polymerase |  |  | 2/195 |  |  |
| 14598 | 246C>T | His82His | T7p29 | gene 5 | DNA polymerase |  |  |  | 4/220 |  |
| 14634 | 282G>A | Leu94Leu | T7p29 | gene 5 | DNA polymerase |  |  |  |  | 4/263 |
| 14734 | 382T>G | Trp128Gly | T7p29 | gene 5 | DNA polymerase |  |  |  | 4/227 |  |
| 14753 | 401A>G | Glu134Gly | T7p29 | gene 5 | DNA polymerase |  |  |  |  | 4/254 |
| 14810 | 458A>G | Glu153Gly | T7p29 | gene 5 | DNA polymerase |  |  |  | 3/237 |  |
| 14849 | 497A>T | Glu166Val | T7p29 | gene 5 | DNA polymerase |  |  | 2/191 |  |  |
| 14866 | 514G>A | Val172Ile | T7p29 | gene 5 | DNA polymerase |  | 22/215 |  |  |  |
| 14896 | 544C>A | Leu182Ile | T7p29 | gene 5 | DNA polymerase |  |  | 2/182 |  |  |
| 14915 | 563A>G | Asp188Gly | T7p29 | gene 5 | DNA polymerase |  | 3/192 |  |  |  |
| 14954 | 602T>C | Val201Ala | T7p29 | gene 5 | DNA polymerase |  |  |  |  | 17/269 |
| 14959 | 607T>G | Tyr203Asp | T7p29 | gene 5 | DNA polymerase |  |  | 2/187 |  |  |
| 14974 | 622T>C | Ser208Pro | T7p29 | gene 5 | DNA polymerase |  |  |  |  | 3/261 |
| 15010 | 658G>T | Ala220Ser | T7p29 | gene 5 | DNA polymerase |  | 2/188 |  |  |  |
| 15057 | 705C>T | Asp235Asp | T7p29 | gene 5 | DNA polymerase |  |  | 3/185 |  |  |
| **15179** | **827A>C** | **His276Pro** | **T7p29** | **gene 5** | **DNA polymerase** | **2/180** |  |  |  |  |
| **15179** | **827A>C** | **His276Pro** | **T7p29** | **gene 5** | **DNA polymerase** | **2/180** |  |  |  |  |
| **15220** | **868A>G** | **Lys290Glu** | **T7p29** | **gene 5** | **DNA polymerase** |  |  |  |  | **3/244** |
| **15316** | **964G>C** | **Ala322Pro** | **T7p29** | **gene 5** | **DNA polymerase** |  | **2/153** |  |  |  |
| **15318** | **966T>C** | **Ala322Ala** | **T7p29** | **gene 5** | **DNA polymerase** |  |  | **4/197** |  |  |
| **15354** | **1002T>C** | **Phe334Phe** | **T7p29** | **gene 5** | **DNA polymerase** |  |  |  | **4/215** |  |
| 15371 | 1019A>G | Asp340Gly | T7p29 | gene 5 | DNA polymerase |  | 3/169 |  |  |  |
| 15410 | 1058C>T | Pro353Leu | T7p29 | gene 5 | DNA polymerase |  | 2/168 |  |  |  |
| 15425 | 1073A>T | Asp358Val | T7p29 | gene 5 | DNA polymerase |  |  | 2/186 |  |  |
| 15449 | 1097A>G | Asp366Gly | T7p29 | gene 5 | DNA polymerase | 209/209 | 168/168 | 175/175 | 188/188 | 225/225 |
| 15461 | 1109A>G | Glu370Gly | T7p29 | gene 5 | DNA polymerase |  |  |  | 2/186 |  |
| 15519 | 1167C>A | Tyr389* | T7p29 | gene 5 | DNA polymerase |  | 2/179 |  |  |  |
| 15569 | 1217G>A | Trp406* | T7p29 | gene 5 | DNA polymerase |  |  | 2/186 |  |  |
| 15633 | 1281G>A | Thr427Thr | T7p29 | gene 5 | DNA polymerase |  |  | 4/196 |  |  |
| 15647 | 1295A>G | His432Arg | T7p29 | gene 5 | DNA polymerase |  |  | 2/187 |  |  |
| 15665 | 1313C>A | Ala438Glu | T7p29 | gene 5 | DNA polymerase |  | 2/189 |  |  |  |
| 15758 | 1406G>A | Trp469* | T7p29 | gene 5 | DNA polymerase | 2/174 |  |  |  |  |
| 15843 | 1491T>C | Ala497Ala | T7p29 | gene 5 | DNA polymerase | 3/135 |  | 2/171 |  | 2/196 |
| 15844 | 1492C>T | His498Tyr | T7p29 | gene 5 | DNA polymerase |  |  | 2/196 |  |  |
| 15845 | 1493A>C | His498Pro | T7p29 | gene 5 | DNA polymerase | 2/155 |  |  |  |  |
| 15848 | 1496A>G | Glu499Gly | T7p29 | gene 5 | DNA polymerase |  | 2/161 | 2/160 |  |  |
| 15850 | 1498A>G | Ile500Val | T7p29 | gene 5 | DNA polymerase |  |  |  | 2/171 |  |
| 15856 | 1504A>T | Asn502Tyr | T7p29 | gene 5 | DNA polymerase |  |  | 5/213 | 7/198 |  |
| 15865 | 1513A>C | Ile505Leu | T7p29 | gene 5 | DNA polymerase |  |  |  | 2/171 |  |
| 15867 | 1515C>T | Ile505Ile | T7p29 | gene 5 | DNA polymerase |  |  |  | 2/198 |  |
| 15871 | 1519A>G | Thr507Ala | T7p29 | gene 5 | DNA polymerase |  |  |  |  | 3/258 |
| 15876 | 1524G>A | Lys508Lys | T7p29 | gene 5 | DNA polymerase | 3/178 |  | 4/208 | 2/194 |  |
| 15879 | 1527C>A | Asn509Lys | T7p29 | gene 5 | DNA polymerase |  |  |  | 3/181 |  |
| 15882 | 1530G>A | Gln510Gln | T7p29 | gene 5 | DNA polymerase |  |  |  | 2/179 |  |
| 15886 | 1534G>A | Ala512Thr | T7p29 | gene 5 | DNA polymerase |  |  | 2/194 |  | 3/246 |
| 15891 | 1539T>C | Ala513Ala | T7p29 | gene 5 | DNA polymerase |  | 3/184 |  |  |  |
| 15893 | 1541A>G | Glu514Gly | T7p29 | gene 5 | DNA polymerase |  |  | 2/190 |  |  |
| 15895 | 1543C>A | Leu515Ile | T7p29 | gene 5 | DNA polymerase | 2/174 |  |  |  |  |
| 15896 | 1544T>C | Leu515Pro | T7p29 | gene 5 | DNA polymerase |  |  | 2/193 |  |  |
| 15908 | 1556A>G | Asp519Gly | T7p29 | gene 5 | DNA polymerase |  |  | 4/197 |  |  |
| 15963 | 1611T>A | Ile537Ile | T7p29 | gene 5 | DNA polymerase |  | 2/181 |  |  |  |
| 16030 | 1678C>T | Pro560Ser | T7p29 | gene 5 | DNA polymerase |  | 2/162 |  |  |  |
| 16060 | 1708C>T | Gln570* | T7p29 | gene 5 | DNA polymerase | 2/165 |  |  |  |  |
| 16099 | 1747G>A | Glu583Lys | T7p29 | gene 5 | DNA polymerase | 2/168 |  |  |  |  |
| 16124 | 1772G>A | Arg591His | T7p29 | gene 5 | DNA polymerase |  |  |  | 2/176 |  |
| 16261 | 1909G>A | Gly637Ser | T7p29 | gene 5 | DNA polymerase |  | 2/197 |  |  |  |
| 16294 | 1942T>C | Tyr648His | T7p29 | gene 5 | DNA polymerase | 2/157 |  |  |  |  |
| 16319 | 1967T>A | Ile656Asn | T7p29 | gene 5 | DNA polymerase | 2/168 |  |  |  |  |
| 16332 | 1980C>T | Cys660Cys | T7p29 | gene 5 | DNA polymerase |  |  |  |  | 5/255 |
| 16416 | 2064T>G | Cys688Trp | T7p29 | gene 5 | DNA polymerase | 2/188 |  |  |  |  |

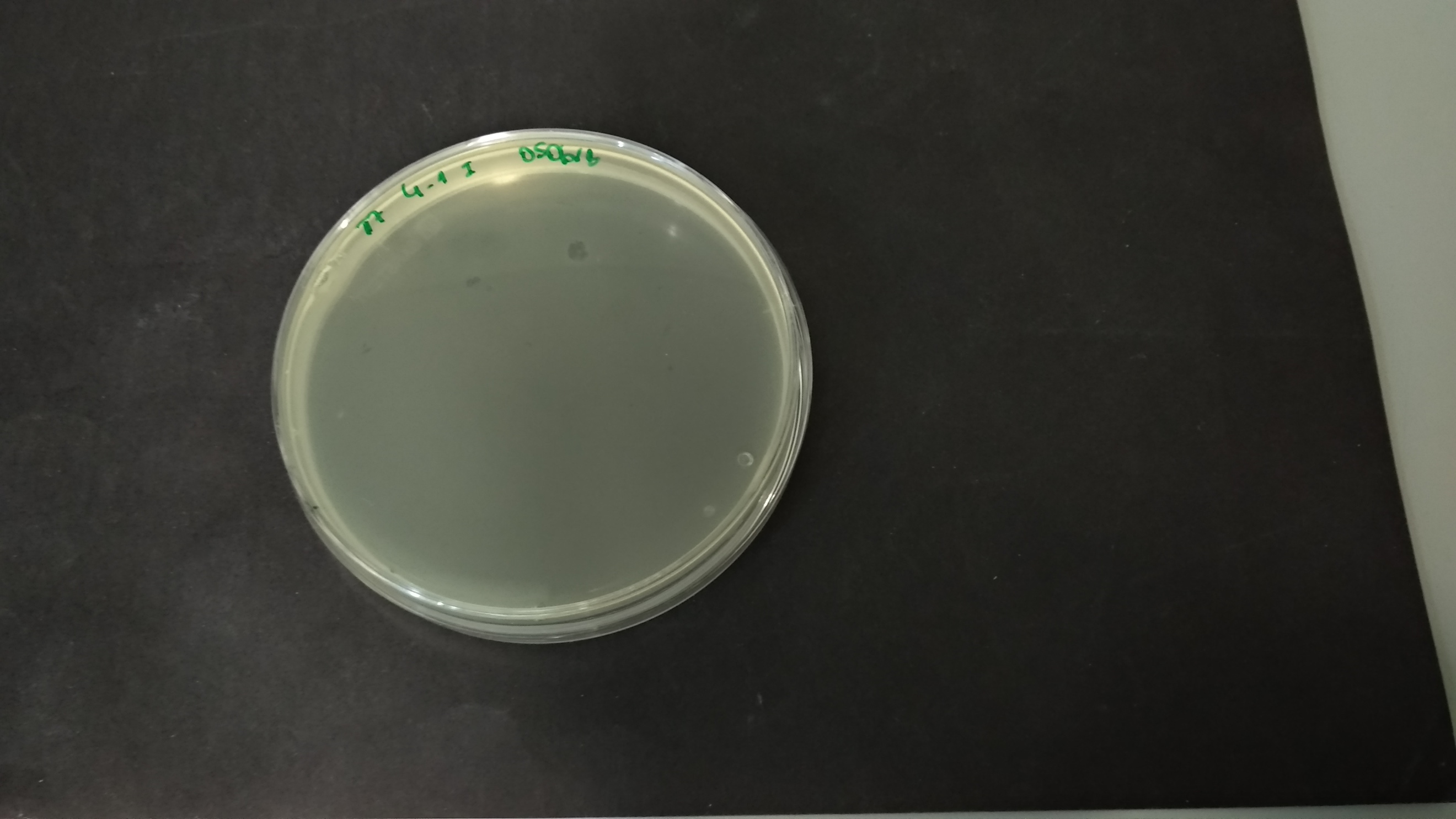

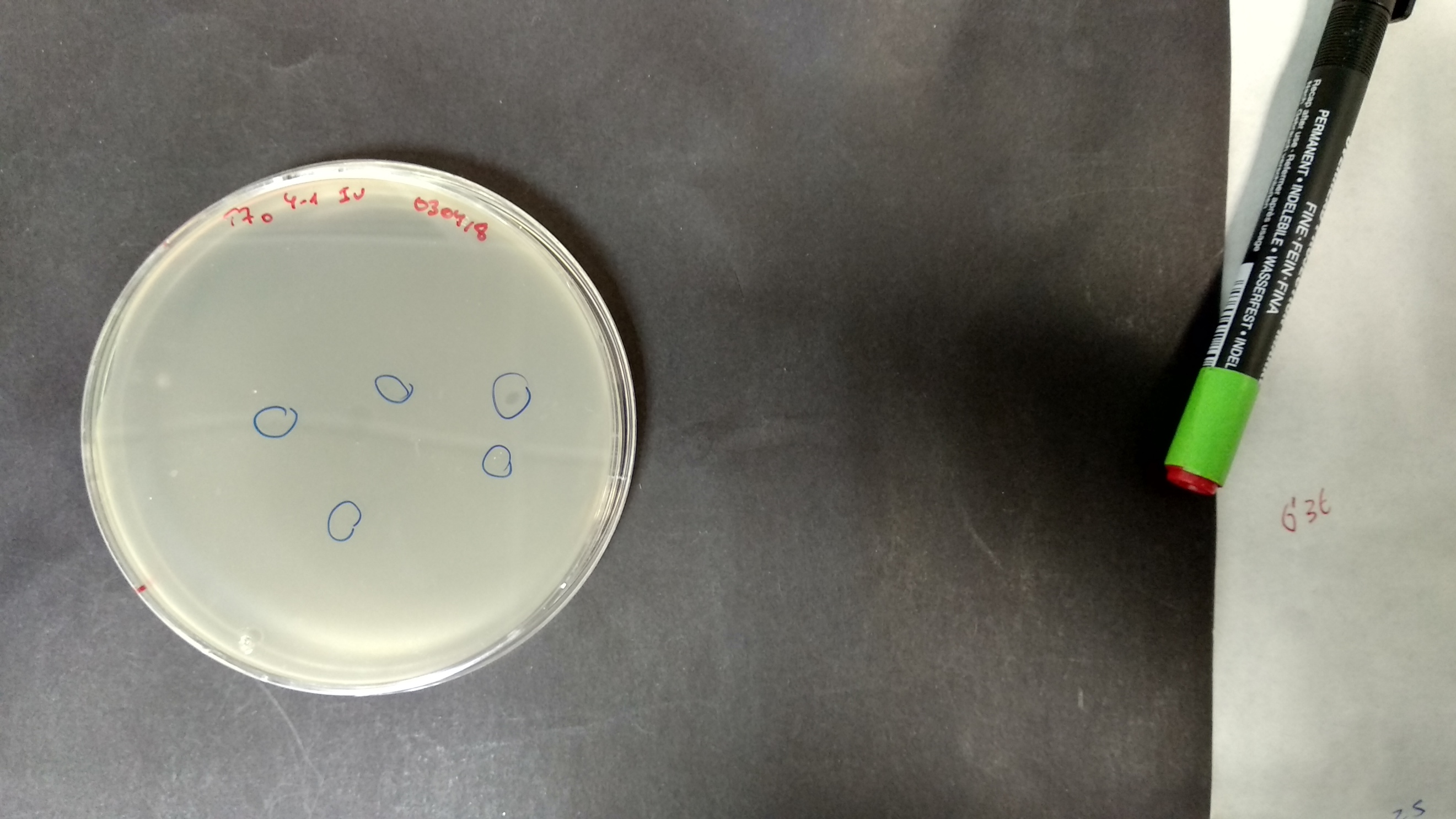

**Fig S1. Pictures of representative plaques corresponding to the initial infection of the engineered *E. coli* strain that contains the alternative LPBCA thioredoxin (the engineered host).**

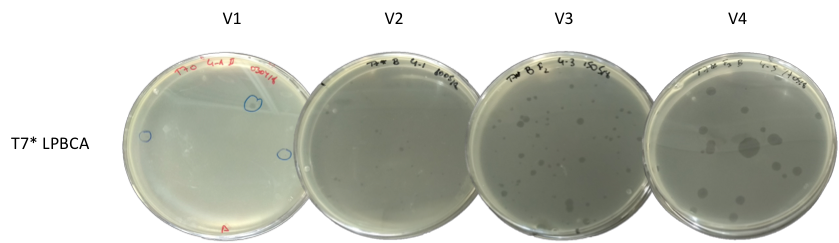

**EXPERIMENT 2 (SANGER)**

**Fig. S2. Pictures showing a representative increase in the size of plaques along the evolution experiment in which the virus adapted to an engineered host containing the alternative LPBCA thioredoxin (engineered host).**

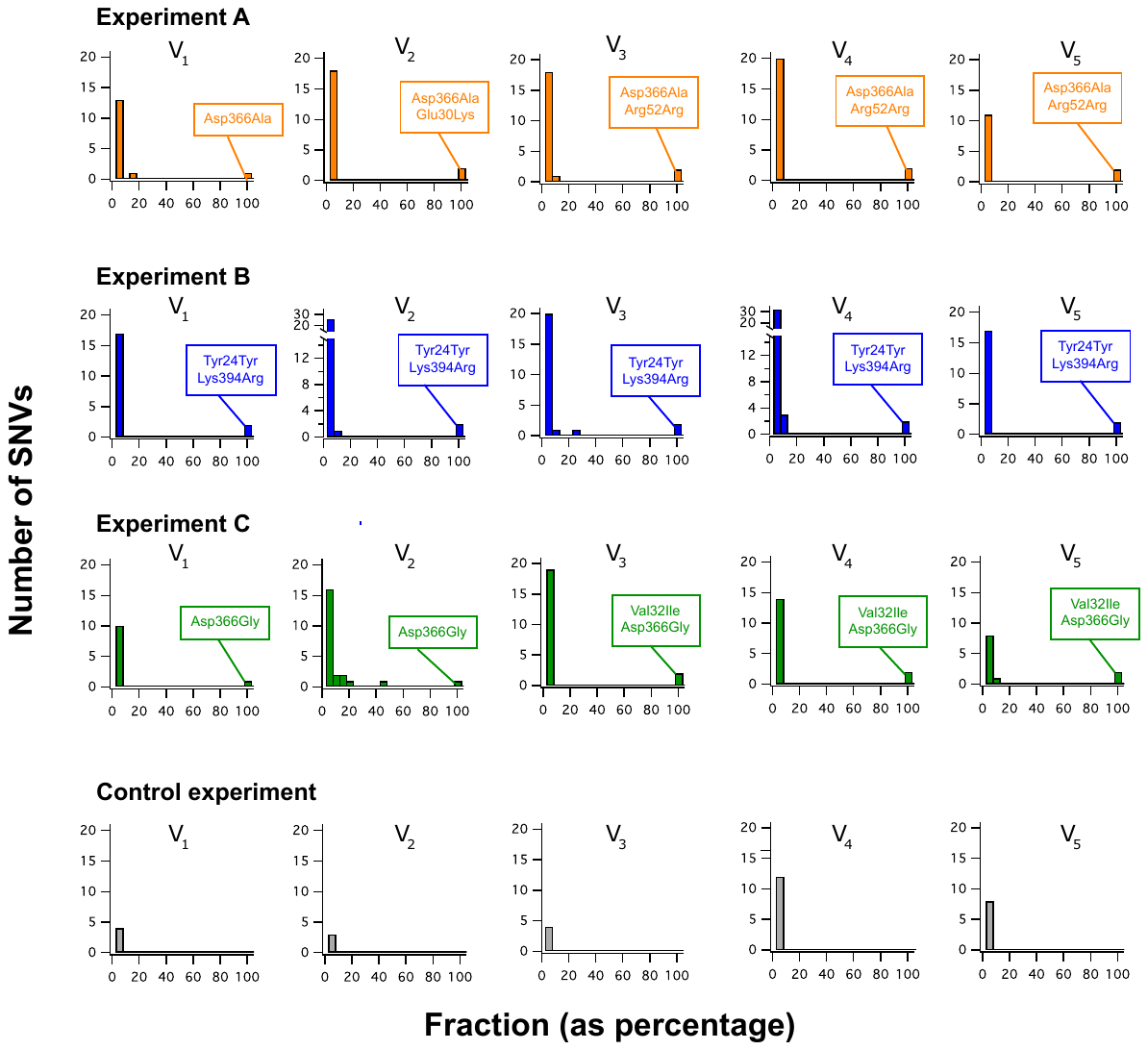

**Fig. S3. Sequence changes in the viral DNA polymerase during laboratory virus evolution. Single nucleotide variants (SNVs) for the gene of the viral DNA polymerase for virus samples from the three evolution experiments of Figure 8. Data were obtained by Illumina sequencing of the DNA extracted after lysis of the engineered host samples. Mutations are binned according to its frequency and the plots show the number of mutations for each 0.05 bin. The mutations that are “fixed” (fraction of occurrence between 0.95 and 1) have been identified. One silent mutation is included because, perhaps, it cannot be ruled out that it could have some effect at the level of RNA structure that translates into some effects on the structure resulting from cotranslational folding. Data for a control experiment corresponding to the propagation of the original virus in the original *E. coli* host is also included.**

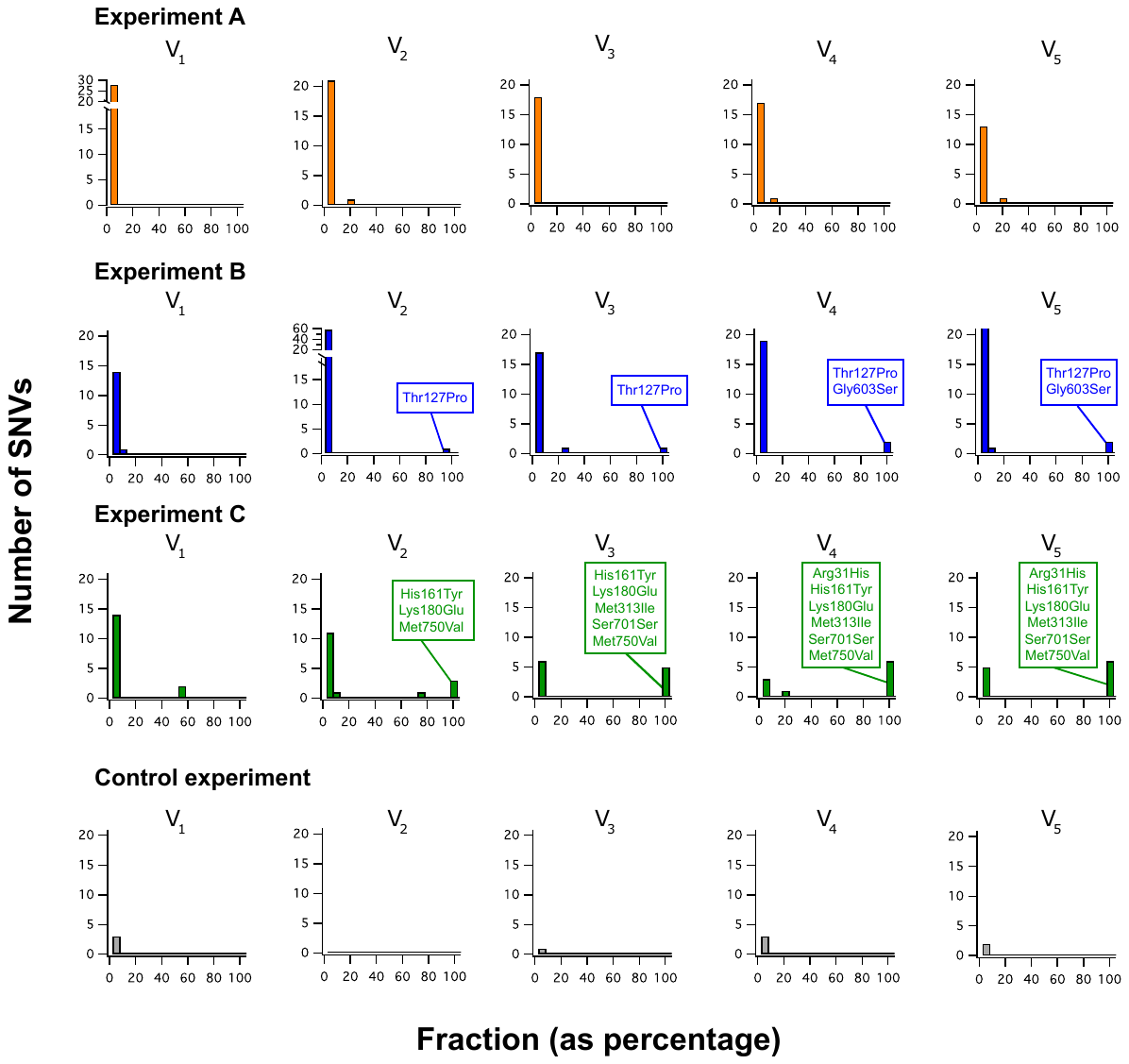

**Fig. S4. Sequence changes in the viral RNA polymerase during laboratory virus evolution. Single nucleotide variants (SNVs) for the gene of the viral RNA polymerase for virus samples from the three evolution experiments of Figure 8. Data were obtained by Illumina sequencing of the DNA extracted after lysis of the engineered host samples. Mutations are binned according to its frequency and the plots show the number of mutations for each 0.05 bin. The mutations that are “fixed” (fraction of occurrence between 0.95 and 1) have been identified. Data for a control experiment corresponding to the propagation of the original virus in the original *E. coli* host is also included.**

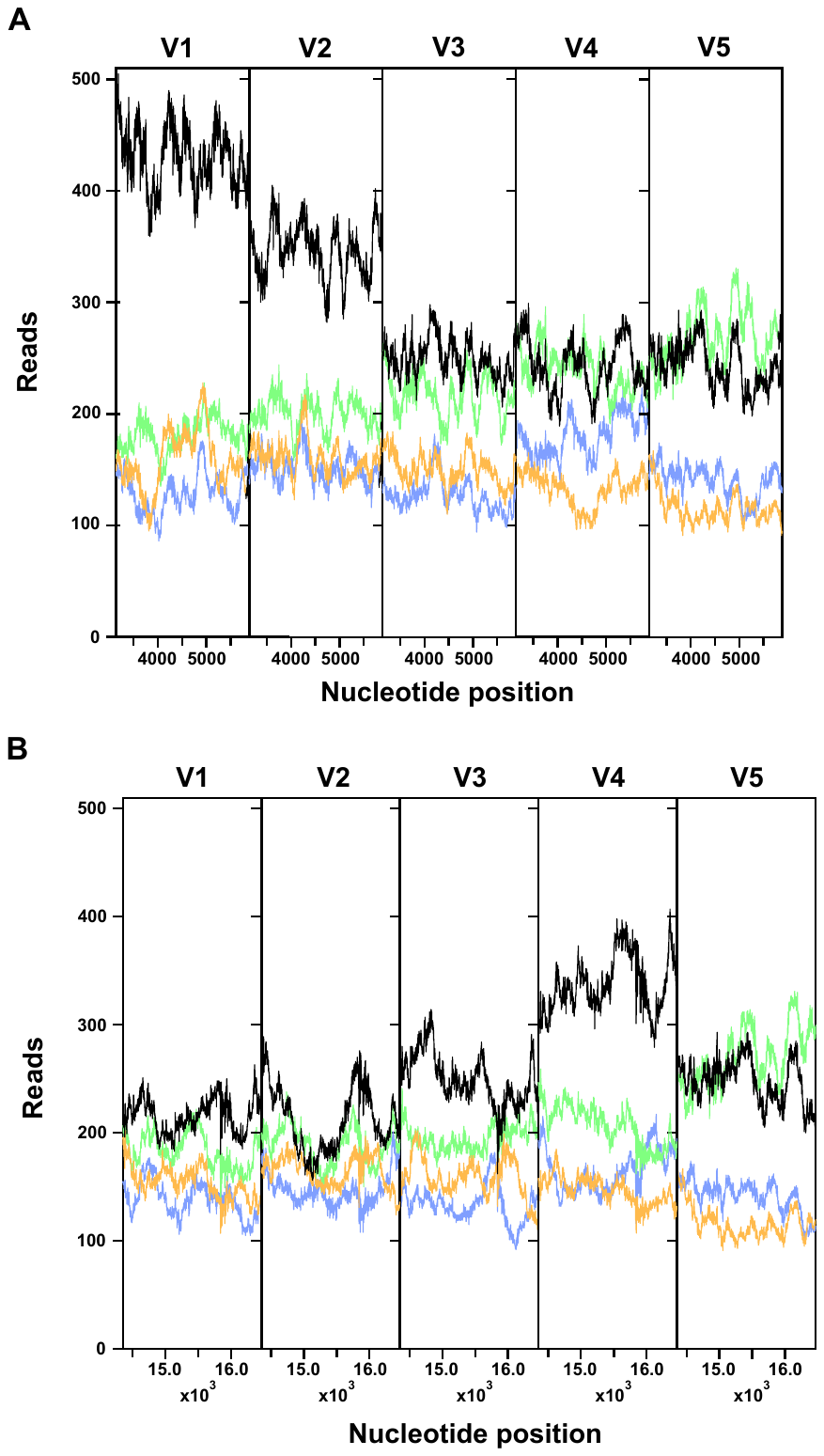

**Fig. S5 Total number of readings for the genes of the viral RNA (A) and DNA (B) polymerases determined from Illumina sequencing of samples from the different rounds of the evolution experiment of the {A,B,C} set. Data for control experiments mimicking the evolution experiment but using the original virus (V_0_) in the original host are also shown.**

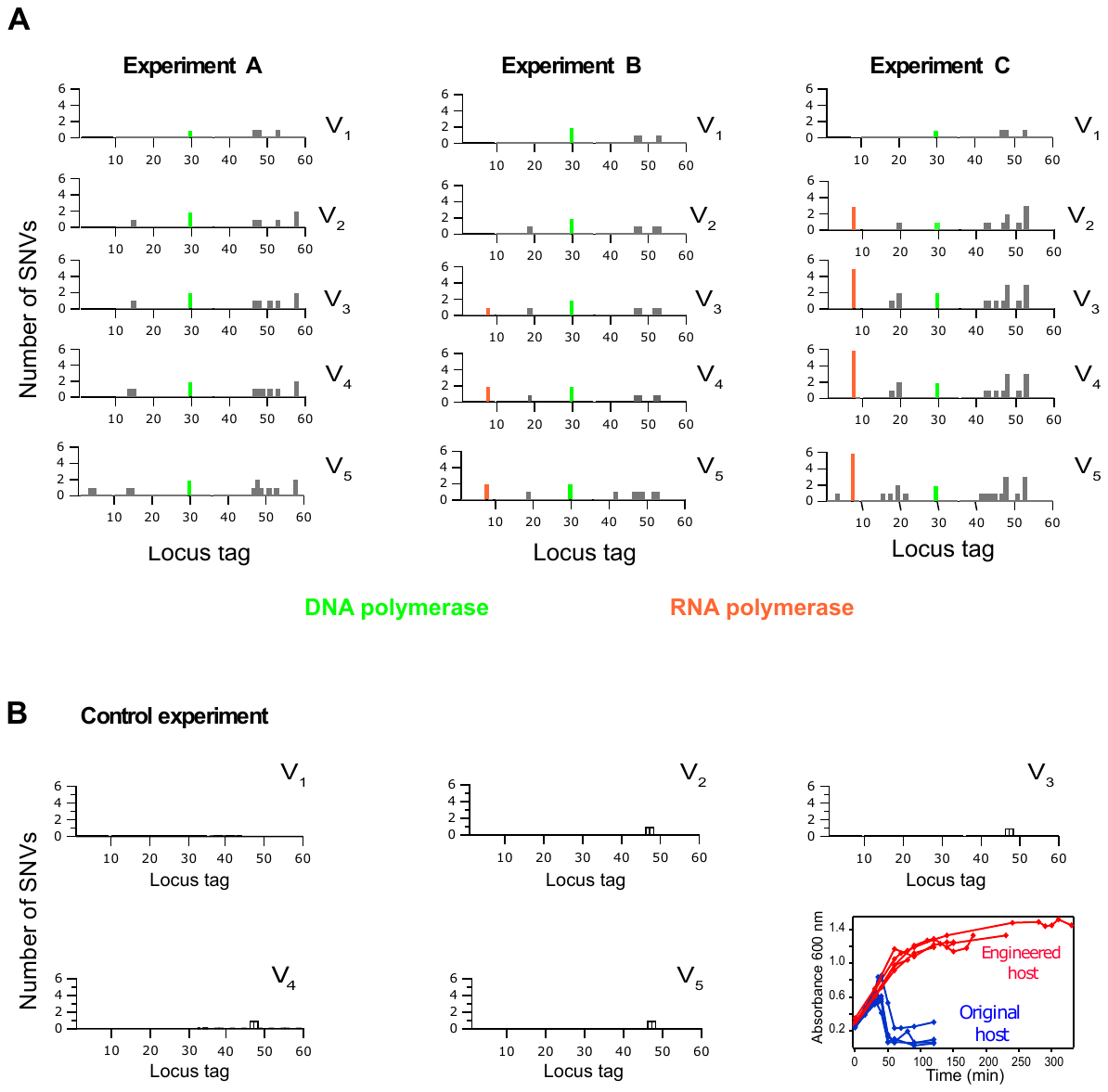

**Fig. S6. Sequence changes in the viral genome during laboratory evolution. a, Number of high frequency (fraction>0.95) mutations for the different viral genes during the evolution experiments of the {A,B,C} set. The viral DNA and RNA polymerases have been highlighted. Data were obtained by Illumina sequencing of the DNA extracted after lysing of the engineered host samples. b, Same as in a, but for a control experiment in which the non-evolved phage propagates in the original host. Black is used here to highlight that this is a control experiment. A panel with lysis curves for this control is included. No selection for propagation in the engineered host is applied in this control experiment and the phage does not evolve the capability to lyse the engineered host.**

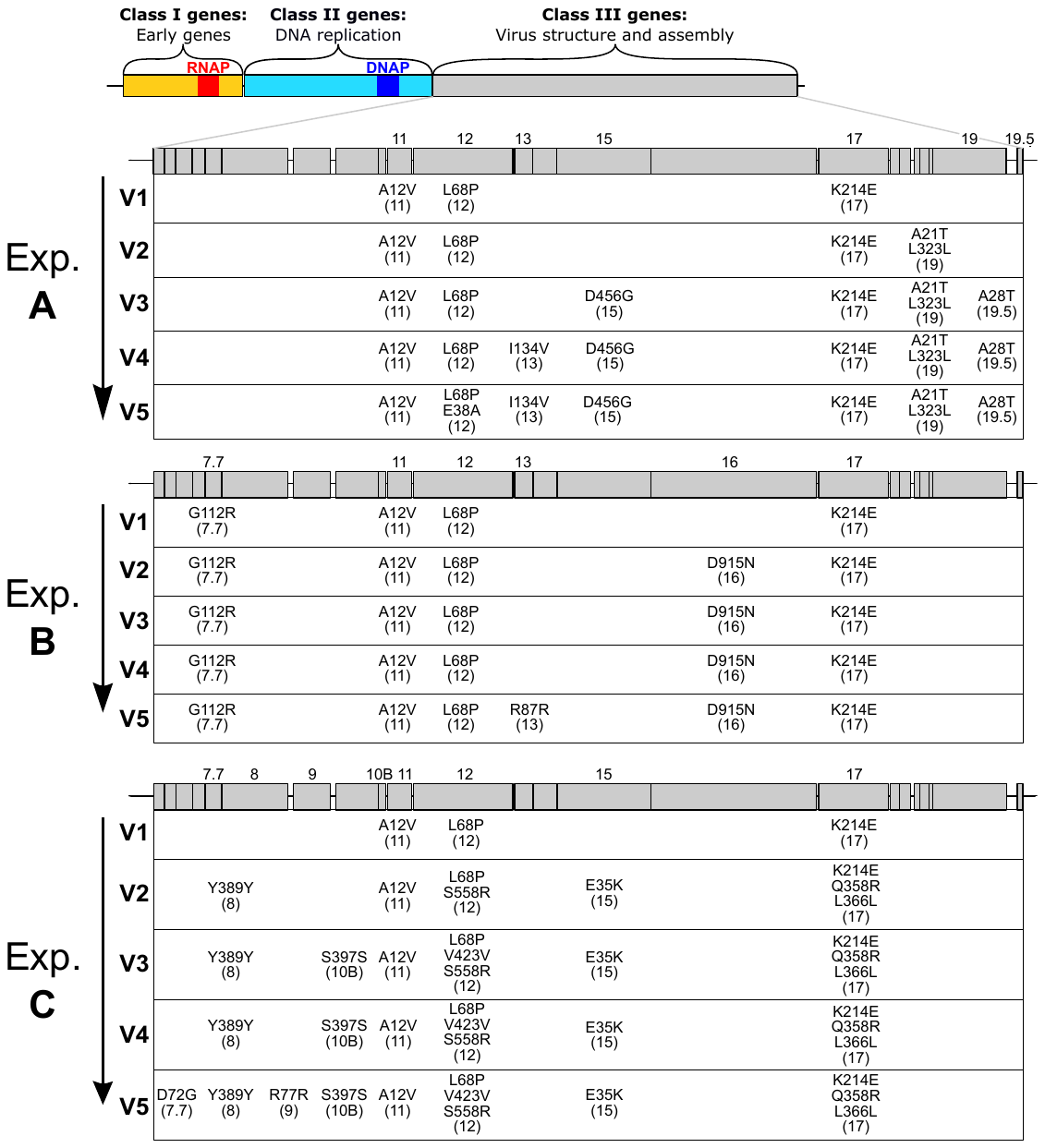

**Fig. S7. Sequence changes in the viral genome during laboratory evolution. a, Number of high frequency (fraction>0.95) mutations for the different viral genes during the evolution experiments of the {A,B,C} set. Only mutations in class III genes are shown here. See figure 9 in the main text for mutations in class I and class II genes. Data were obtained by Illumina sequencing of the DNA extracted after lysing of the engineered host samples (see main text and Figure 7 for details). A control experiment in which the non-evolved virus propagates in the original host yielded very few mutations in class III genes (see Figure S6).**
